# Sentinel Plants Enable Aboveground Detection of Belowground Soil Microbial Activity

**DOI:** 10.64898/2026.03.23.713735

**Authors:** Yunqing Wang, John P. Marken, Eugene Li, Paul T. Tarr, Elliot M. Meyerowitz, Gozde S. Demirer

## Abstract

Rhizosphere microbial processes play a central role in soil function and plant health yet remain difficult to monitor noninvasively. Engineered sentinel plants that use bacterial-to-plant communication channels are promising. However, no such efforts have thus far enabled a detectable aboveground response in the sentinel plant. Here, we optimize a previously described synthetic bacteria-to-plant communication channel based on the *p*-coumaroyl-homoserine lactone (pC-HSL) signaling molecule in plants to function as aboveground sentinels of belowground microbial activities. *Arabidopsis thaliana* sentinel plants harboring this optimized circuit detect root-applied pC-HSL at concentrations as low as 30 nM in roots and 3 μM in leaves, demonstrating long-distance signal transmission from below ground to aboveground tissues. Moreover, sentinel plants report pC-HSL production by engineered *Escherichia coli* and *Pseudomonas putida* colonizing plant roots in both plate and soil assays. These results establish an engineered plant platform that converts rhizosphere microbial activity into a visible aboveground signal, enabling a minimally invasive platform for monitoring rhizosphere microbial gene expression and for precision agriculture and soil management.

## Introduction

The rhizosphere, a narrow zone of soil directly influenced by plant roots, is an essential component of soil health and agricultural sustainability. Rhizosphere microbial communities drive nutrient cycling, affect greenhouse gas emissions, and shape crop productivity^1^. Despite this key ecological importance, microbial activity in the rhizosphere remains difficult to monitor because existing methods rely on destructive sampling or indirect chemical measurements that lack sufficient spatiotemporal resolution^2^. Sensing approaches that do not perturb and can resolve underground microbial activity could advance the study and management of soil health for global sustainability^3^.

Among emerging strategies for rhizosphere monitoring, microbial biosensors couple soil inputs to quantifiable outputs. These systems consist of genetically engineered bacteria that integrate input-responsive transcriptional regulators with optical or electrochemical reporters, thereby forming programmable input-output circuits^2^. Microbial environmental biosensors have been developed to detect nutrient availability and pollutants, such as heavy metals and organic contaminants^4–6^. Despite their versatility, deployment of microbial biosensors in open soil environments is constrained by challenges of containment and regulatory uncertainty. They also rely on specialized instrumentation for signal readout. Their spatial sensing is also inherently limited because microbes sample only small, heterogeneous soil microenvironments, though recent advances in hyperspectral imaging offer a path toward scalable, field-level detection^7^.

Plants offer an alternative chassis for environmental biosensing. Unlike microbial biosensors, plants inherently integrate and transmit signals from the rhizosphere to aboveground tissues. They are self-powered via photosynthesis, and their capacity to generate diverse outputs, including pigments, scents, and volatile organic compounds, can enable detection at scale without specialized instrumentation^8^. Plants have been engineered to report a wide range of stressors, including osmotic stress, γ-radiation, insecticides, and contaminants^9,10^. One example is the detection of the explosive trinitrotoluene via a visible de-greening response^11^. Another study reprogrammed the abscisic acid receptor to detect non-native ligands, such as fungicides and insecticides, achieving nanomolar sensitivity with outputs spanning stomatal closure, thermal signatures, and pigments^12^. Beyond abiotic stress, sentinel plants have been developed to detect biotic stressors. Field trials with engineered tobacco expressing a fluorescent protein under the control of synthetic salicylic acid-responsive promoters provided early warnings of pathogen infection^13^. In parallel, commercial efforts have begun to translate these concepts into agricultural applications. InnerPlant created engineered soybean plants that fluoresce upon fungal infection, and Insignum developed maize that accumulates anthocyanins in response to fungal challenge^14,15^.

Building on these advances, recent work has explored plants’ ability to sense microbially produced signaling molecules. Boo et al. established a cross-kingdom signaling system by engineering *Pseudomonas putida* and *Klebsiella pneumoniae* that produce *p*-coumaroyl-homoserine lactone (pC-HSL), which was detected by a synthetic circuit in plant roots^16^. This work demonstrated that plants can be engineered to respond to microbially synthesized signals, but detection was limited to root-localized responses. Therefore, it remains to be shown whether these microbial signals can be transmitted to and detected in aboveground plant tissues, a requirement for scalable, noninvasive monitoring in realistic soil environments.

We engineered sentinel plants that convert rhizosphere microbial activity into a visible aboveground signal using pC-HSL as a cross-kingdom communication molecule. Capitalizing on pC-HSL’s unique advantages, including its orthogonality to endogenous plant and soil signaling pathways^17^, genetic encodability in diverse microbial hosts^18,19^, and the availability of known receptors, we optimized a pC-HSL-responsive plant circuit and generated *Arabidopsis thaliana* plants that sensitively detect microbially synthesized pC-HSL in both plate and soil environments. This work demonstrates that plants can perceive a bacterially produced signaling molecule and report this information in aboveground tissues, enabling a noninvasive readout of rhizosphere microbial gene expression. More broadly, establishing a direct communication channel between underground microbes and plant leaves introduces a paradigm for facile plant-based biosensing and creates opportunities for continuous, field-deployable monitoring of diverse soil processes.

## Results

### Optimization of the pC-HSL-inducible circuit via transient Nicotiana benthamiana assays

To establish a sentinel plant with high detection sensitivity and dynamic range, we first optimized a pC-HSL-inducible genetic circuit. This involved optimizing both the nuclear localization of the transcriptional activator RpaR, which binds pC-HSL and dimerizes, and the repeat number of its cognate operator sequence RpaO. A library of circuit variants was constructed by systematically altering the number of RpaO operator sites (0x, 2x, 4x, or 6x) upstream of the reporter gene and by testing alternative nuclear localization signals (NLSs) (SV40 or bipartite BP) to modulate the strength of RpaR-VP16 nuclear import (**Fig. 1A**)^20^. To account for batch-to-batch and plant-to-plant variability in the optimization process, reporter activity was quantified using a dual-luciferase assay, in which Green-enhanced Nano-lantern (GeNL) served as the inducible reporter normalized to a constitutively expressed NanoLuc (Nluc), measured with a plate reader^21,22^. Each construct was expressed in *N. benthamiana* leaves via *Agrobacterium*-mediated infiltration, and the leaves were treated with 0 to 10 μM pC-HSL after 48 h (**Supplementary Fig. 1A**).

**Figure 1.**
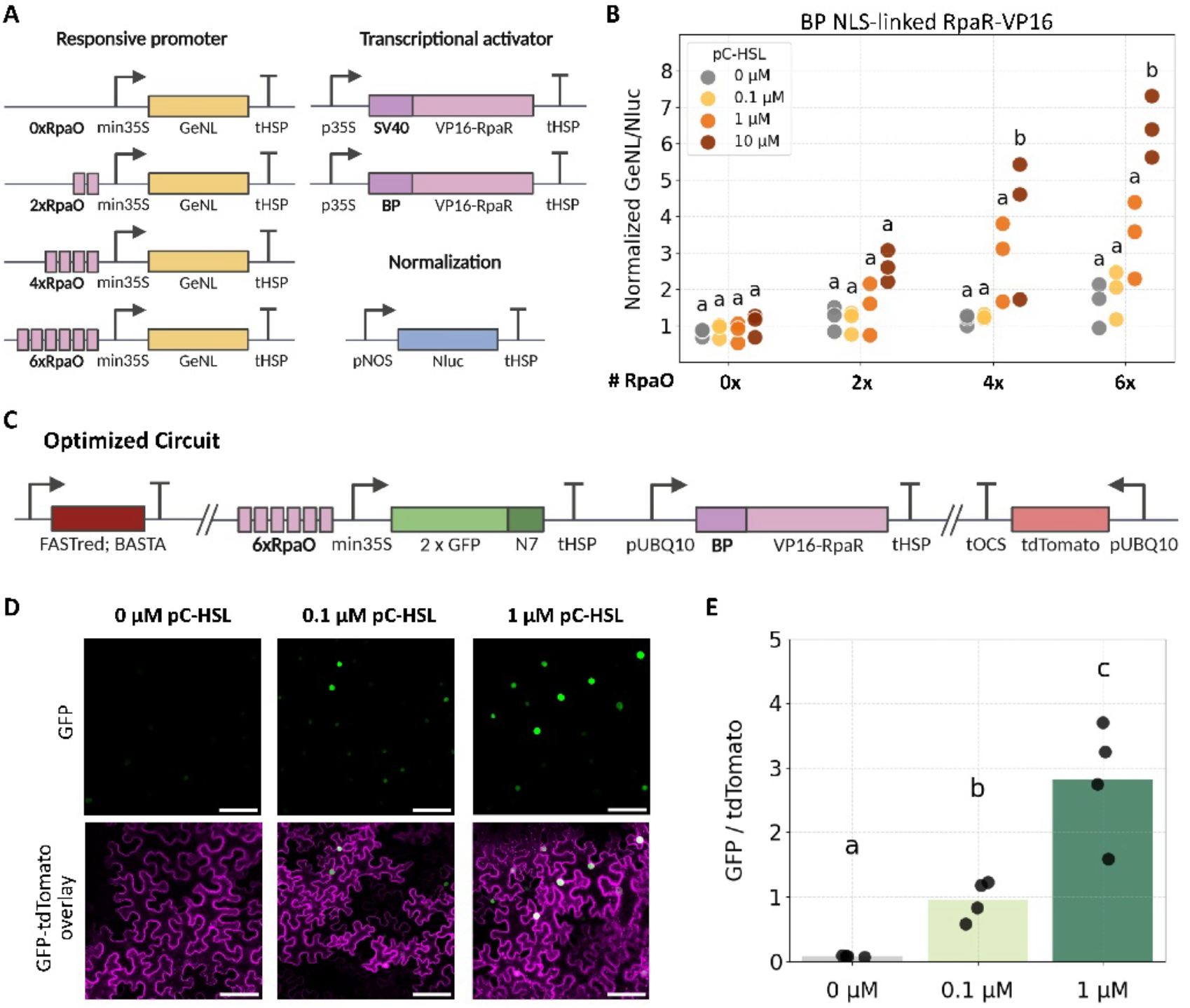
Optimization of pC-HSL-inducible synthetic circuits in *N. benthamiana* transient assays. **A)** Circuit design for combinations of RpaO operator site numbers and nuclear localization signal (NLS). **B)** Normalized GeNL/ NanoLuc activity of circuit variants treated with 0, 0.1, 1, or 10 µM pC-HSL. n = 3 biological replicates. **C)** The optimized pC-HSL-inducible circuit used for fluorescence assays contains 6×RpaO-mini35S-2×GFP-N7-tHSP inducible output module, pUBQ10-BP-VP16-RpaR-tHSP transcriptional activator, pUBQ10-tdTomato-tOCS normalization module in the reverse orientation, and FAST-Red and BASTA selection markers. **D)** Representative confocal microscopy images of GFP and GFP/tdTomato overlay in *N. benthamiana* leaves transiently expressing the optimized pC-HSL-inducible circuit treated with 0, 0.1, or 1 μM pC-HSL. Scale bars, 100 μm. **E)** Quantification of sensor response as GFP normalized to tdTomato fluorescence across treatments. Bars represent mean ± S.E.M. of 4 biological replicates. Log-transformed data were analyzed by one-way ANOVA with Tukey’s HSD post hoc test. Different letters indicate statistically significant differences (p < 0.05).

Circuits containing six operator sites upstream of a minimal 35S promoter exhibited the largest fold change to pC-HSL (**Fig. 1B**). Dose-response analysis of the BP NLS-linked transcriptional activator design revealed induction ratios of 1.1-fold at 0.1 μM, 2.5-fold at 1 μM, and 3.7-fold at 10 μM pC-HSL (**Supplementary Table 1**). Induction ratios for the SV40 NLS variants were similar (**Supplementary Fig. 1B**), but we selected the BP NLS design for subsequent work because it displayed lower variability across biological replicates. These results establish that both operator multiplicity and nuclear import affect transcriptional response in plant synthetic circuits.

We next adapted the optimized circuit configuration to enable fluorescence output for studying spatial response patterns in plant tissues. To this end, we replaced GeNL with a nuclear-localized 2×GFP and replaced NanoLuc with a constitutively expressed membrane-localized tdTomato. Seed coat marker FAST-Red and BASTA antibiotic resistance cassettes were also incorporated for subsequent use of this construct to generate transgenic lines (**Fig. 1C**)^23^. Transient *N. benthamiana* leaf assay of the optimized GFP circuit via confocal microscopy confirmed strong nuclear-localized fluorescence in response to 0.1 and 1 μM pC-HSL, while untreated leaves displayed minimal signal (**Fig. 1D**). Confocal microscopy images from 3 additional biological replicates containing individual plants are provided in **Supplementary Fig. 2**. Quantification of confocal images across all 4 biological replicates showed a robust, dose-dependent response detectable at as low as 0.1 µM pC-HSL (**Fig. 1E**). The circuit exhibited an 8.2-fold increase at 0.1 μM and a 36-fold increase at 1 μM relative to the untreated control (**Fig. 1E** and **Supplementary Table 2**). These results are consistent with the plate reader-based luciferase assay, with the larger fold changes reflecting the higher sensitivity of fluorescence-based imaging readouts.

### pC-HSL sensing in the roots of Arabidopsis thaliana sentinel plants

We next generated stably transformed *A. thaliana* lines carrying the optimized pC-HSL-inducible GFP circuit (**Fig. 1C**) to evaluate root responses to exogenous pC-HSL *in vivo*. Seedlings were treated with pC-HSL, and roots were imaged using confocal microscopy to analyze activation of the inducible circuit (**Supplementary Fig. 3A**). Three independent transgenic lines were generated and screened for reporter responsiveness to pC-HSL. The best performing stable line (Line #1) was selected for further characterization and experiments (**Supplementary Fig. 4-6**).

GFP fluorescence was detected in all roots in response to pC-HSL treatment, while untreated controls exhibited little to no signal (**Fig. 2A**). Quantitative analysis of confocal images across nine biological replicates, collected in two independent experiments, showed that the circuit remained fully functional in root tissues, displaying a detection limit of 0.03 µM pC-HSL and a gradual fluorescence intensity increase up to 1 µM pC-HSL (**Fig. 2B**). Induction at 30 nM pC-HSL was 25-fold compared to the untreated control. The magnitude of induction overall was significantly higher in the stable lines compared to the transient *N. benthamiana* assays: transient assays showed an 8.2-fold increase at 0.1 μM and a 36-fold increase at 1 μM, whereas stable line #1 showed a 61-fold increase at 0.1 μM and a 2600-fold increase at 1 μM relative to the untreated control (**Supplementary Table 3**).

**Figure 2.**
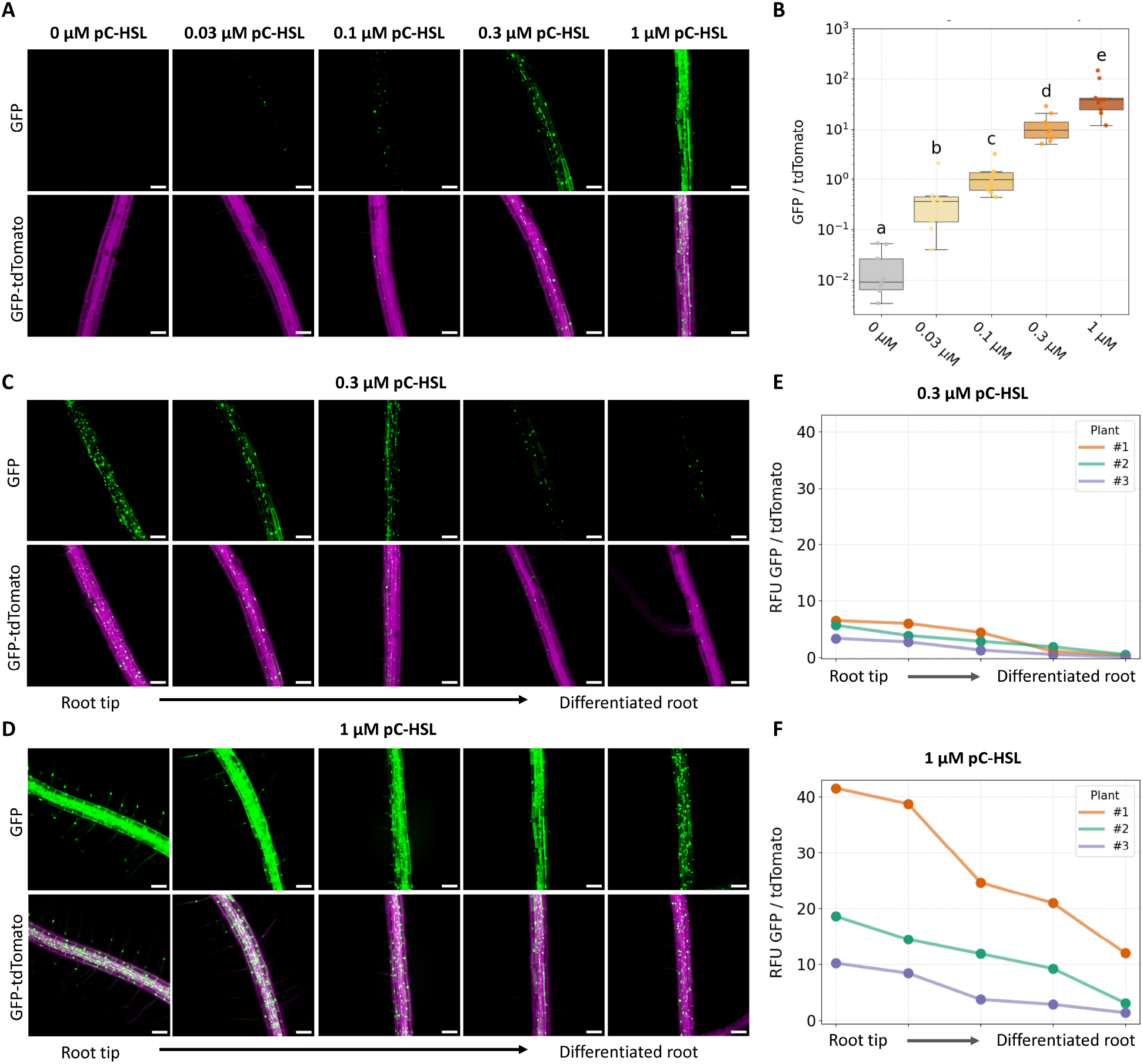
Characterization of pC-HSL-inducible circuit in roots of stably transformed *A. thaliana*. **A)** Representative confocal microscopy images of GFP (green) and tdTomato (magenta) signals in *A. thaliana* roots treated with pC-HSL at concentrations ranging from 0 to 1 μM. **B)** Quantification of circuit activation in roots of stably transformed *A. thaliana* seedlings from confocal images. GFP fluorescence was measured and normalized to tdTomato in the primary root. Data were shown as boxplots (median, Interquartile Range IQR, and 1.5×IQR whiskers) with individual data points overlaid (n = 9 biological replicates). Log-transformed data were analyzed by one-way ANOVA with Tukey’s HSD post hoc test. Different letters indicate statistically significant differences (p<0.05). **C, D)** Representative images of the spatial profile of GFP activation along the primary root treated with 1 μM and 0.3 μM pC-HSL. Images are shown from bottom to top as fields of view progressing from the root tip toward the maturation zone. **E, F)** Quantification of circuit activation along the primary roots when treated with 1 μM and 0.3 μM pC-HSL. Each line represents an individual biological replicate. Scale bars, 100 μm.

Notably, the GFP reporter activation was not spatially uniform along the sentinel plant root longitudinal axis. GFP fluorescence was strongest in the root tip and decreased toward the root elongation and maturation regions, as observed in representative confocal microscopy images following treatment with 1 µM or 0.3 µM pC-HSL (**Fig. 2C, D**, and **Supplementary Fig. 7**). Quantification of reporter activation from the confocal microscopy images along the length of individual primary roots across three independent biological replicates confirmed the gradient of GFP response to pC-HSL decreasing from root tip to top region (**Fig. 2E, F**).

In addition to the longitudinal GFP analysis, we also analyzed the radial response pattern of sentinel roots to pC-HSL treatment. A reconstructed transverse section showed reporter activation across all root cell layers at the highest dose level, 1 µM. GFP fluorescence was most strongly and frequently detected in epidermal and cortical layers and extended toward the endodermis and stele. At lower pC-HSL concentrations of 0.3 µM, little to no activation was observed in the inner endodermis and stele layers, whereas at higher pC-HSL concentrations of 1 µM, reporter activation was frequently detected in all root cell layers (**Supplementary Fig. 8**).

### pC-HSL sensing in the leaves of Arabidopsis thaliana sentinel plants

Having established a sensitive response to pC-HSL in sentinel plant roots, we next investigated whether pC-HSL can travel acropetally to induce a response in leaf tissue, a key property for plants capable of visibly reporting underground rhizosphere activity. Sentinel *A. thaliana* lines harboring the optimized circuit were grown on split MS agar plates, in which pC-HSL was applied exclusively to the lower half containing the root system, and the response was measured in leaves after 72 h (**Supplementary Fig. 3B**).

Confocal microscopy imaging showed the sensor’s fluorescence response in leaf tissues at concentrations as low as 1 µM of exogenously applied pC-HSL to the roots (**Fig. 3A**). At higher pC-HSL concentrations of 3 and 10 µM, GFP fluorescence in leaves was significantly higher (**Fig. 3A**). Quantitative analysis of confocal microscopy images across nine biological replicates, collected in two independent experiments, revealed that the GFP intensity increased with the concentration of pC-HSL applied to the roots (**Fig. 3B**). For each leaf sample, three fields of view (FOVs) were captured, and the integrated fluorescence densities from the entire FOVs were averaged to obtain representative measurements (**Supplementary Fig. 9**). The circuit in the leaf exhibited a 2.1-fold increase at 1 μM, a 270-fold increase at 3 μM, and an 840-fold increase at 10 μM relative to the untreated control (**Supplementary Table 4**).

**Figure 3.**
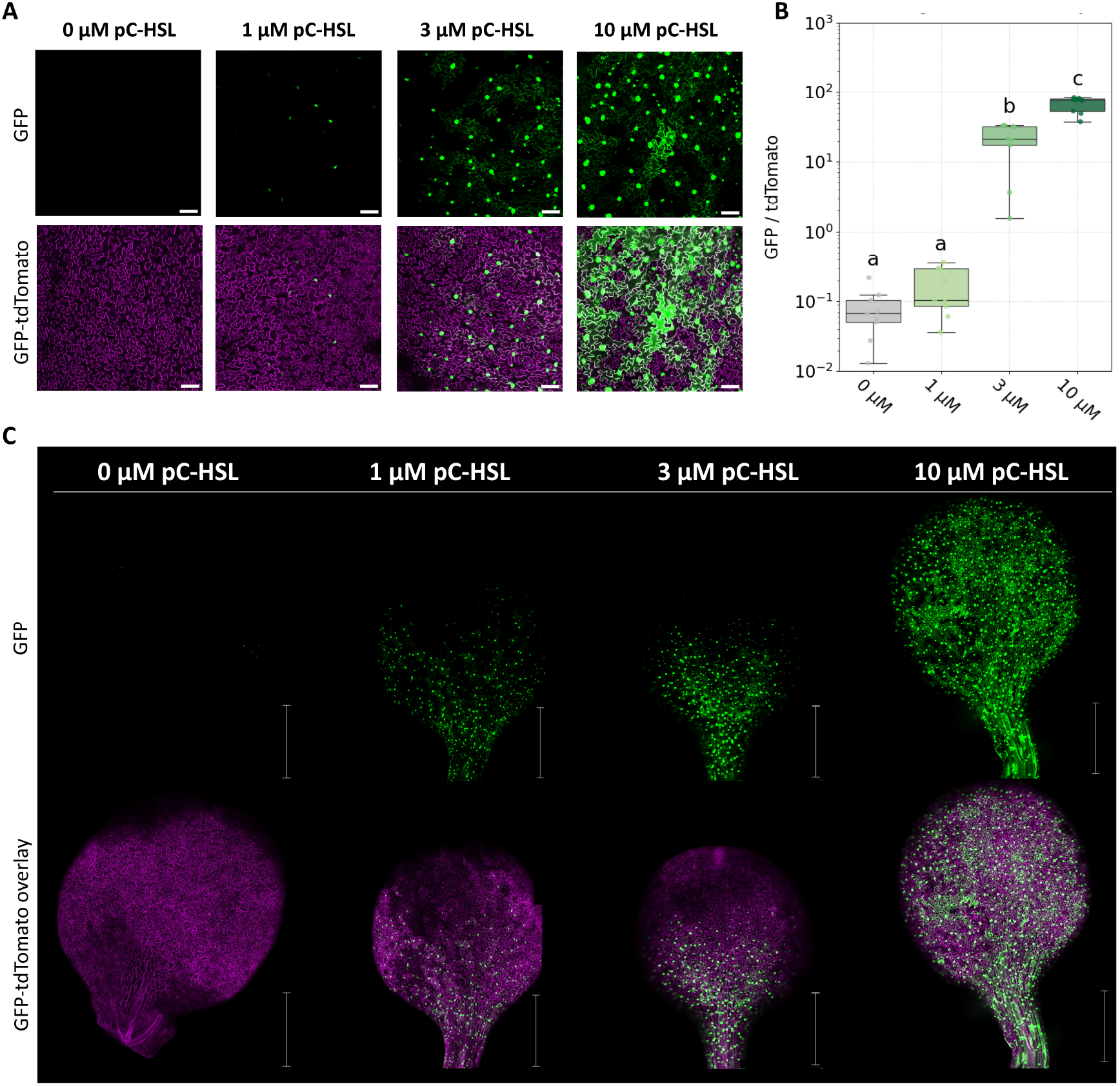
Characterization of pC-HSL-inducible circuit in leaves of stably transformed *A. thaliana* plants. **A)** Representative confocal images of GFP (green) and tdTomato (magenta) signals in *A. thaliana* leaves with root-treated pC-HSL at concentrations ranging from 0 to 10 μM. Scale bars, 100 μm. **B)** Quantification of circuit activation in leaves of stably transformed *A. thaliana* seedlings. Seedlings were grown on split MS agar plates in which pC-HSL was applied only to the lower region. Data were shown as boxplots (median, Interquartile Range IQR, and 1.5×IQR whiskers) with individual data points overlaid (n = 9 biological replicates). Log-transformed data were analyzed by one-way ANOVA with Tukey’s HSD post hoc test. Different letters indicate statistically significant differences (p<0.05). **C)** Whole-leaf confocal scan of systemic activation. GFP fluorescence is spatially patterned, appearing strongest near basal areas of the leaf. Scale bars, 1 mm.

Whole-leaf confocal microscopy imaging of root-treated plants revealed the spatial pattern of sensor response in leaves, showing that the reporter fluorescence was concentrated and stronger in leaf areas near the petiole (**Fig. 3C**). Consistent with this spatial pattern, GFP fluorescence within individual leaves was predominantly localized to areas close to vasculature and veins, with a stronger signal observed in vascular-associated regions (FOV1) compared to non-vascular regions (FOV2 and FOV3) when lower pC-HSL concentrations of 1 and 3 μM were used. At a high pC-HSL concentration of 10 μM, the sensor response is readily detected across all parts of the leaf (**Supplementary Fig. 9**).

In addition to this spatial expression pattern, circuit responsiveness also varied with leaf developmental stage. Cotyledons showed stronger circuit activation and brighter GFP compared to the newly emerging younger true leaves, both in areas near and further away from the vasculature and veins (**Supplementary Fig. 10**). Young true leaves also displayed lower constitutive tdTomato marker signals, indicating that weaker circuit expression might explain the reduced activation levels by pC-HSL in young leaves (**Supplementary Fig. 10**).

### Sentinel plants detect bacterially produced pC-HSL in root-microbe co-culture systems

The rhizosphere is a chemically dynamic environment where microbial communities continuously exchange signals with each other and the surrounding plants^24^. To move beyond exogenous chemical induction and toward a system that can directly report native microbial activity, we tested whether pC-HSL produced by engineered bacteria could activate the sentinel plant response. Establishing this capability would allow microbial activity to be encoded as a chemical signal that the sentinel plant detects and reports.

To this end, we reconstructed the pC-HSL biosynthetic pathway in *Escherichia coli* and *Pseudomonas putida*. The synthetic biosynthetic pathway consisted of three enzymes: TAL from *Rhodobacter sphaeroides*, 4CL from *Nicotiana tabacum*, and the pC-HSL synthase RpaI from *Rhodopseudomonas palustris*, which convert L-tyrosine into p-coumarate, p-coumaroyl-CoA, and ultimately pC-HSL, respectively (**Supplementary Fig. 11A**)^18^. The synthetic operon expressed these enzymes with a constitutive promoter and a strong intrinsic terminator (**Supplementary Fig. 11B**). Both engineered bacterial strains produced minimal pC-HSL in the absence of the precursor and accumulated substantially higher levels in supernatant when supplemented with the precursor p-coumarate, and measured by a fluorescence bioassay (see Methods). At 48 h, *E. coli* produced 32.8 µM pC-HSL in the bacterial supernatant when cultured with the precursor p-coumarate and 3.0 µM without, while *P. putida* produced 13.9 µM and 2.1 µM with and without p-coumarate, respectively (**Supplementary Fig. 11C-D**).

We next evaluated whether microbially generated pC-HSL could trigger a detectable response *in planta* using a root-microbe co-culture assay. Sentinel *Arabidopsis thaliana* seedlings were grown on MS agar plates containing p-coumarate, and bacterial suspensions were applied exclusively to the lower half of the plate to ensure contact only with the root system. After three days of co-cultivation, GFP fluorescence was measured in leaves (**Supplementary Fig. 3C**). Representative confocal images of leaf tissues showed that both engineered *E. coli* and *P. putida* induced strong reporter activation in leaves when roots were inoculated with as low as OD 1 bacteria, with higher bacterial OD leading to progressively stronger GFP signal (**Fig. 4A and C**). Quantitative analysis of confocal microscopy images across six biological replicates showed that, relative to uninoculated controls, *E. coli* induced a 15-fold increase in reporter signal at OD 1, a 44-fold increase in reporter signal at OD 3, and a 140-fold increase at OD 10. Similarly, *P. putida* induced a 12-fold increase in reporter signal at OD 1, a 46-fold increase at OD 3, and a 200-fold increase at OD 10 (**Fig. 4B and D, Supplementary Table 5**).

**Figure 4.**
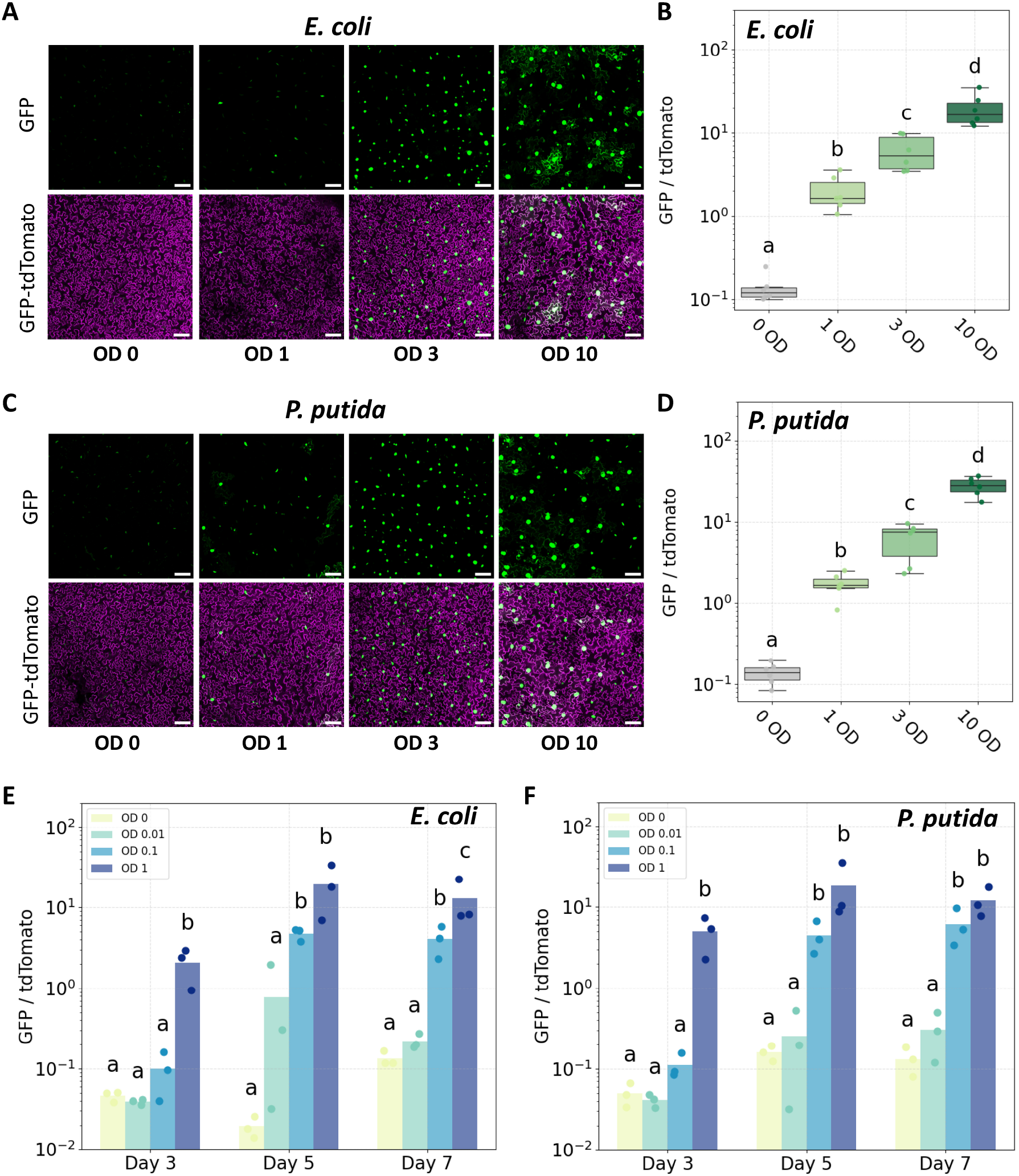
Sentinel plants detect bacterially secreted pC-HSL in co-culture systems. **A, C)** Representative confocal images of GFP (green) and tdTomato (magenta) signals in *A. thaliana* leaves with root-treated *E. coli* (top) or *P. putida* (bottom) suspension at OD600 ranging from 0 to 10. **B, D)** Quantification of circuit activation in *A. thaliana* leaves with root-treated *E. coli* (top) or *P. putida* (bottom) suspension. Data were shown as boxplots (median, Interquartile Range IQR, and 1.5×IQR whiskers) with individual data points overlaid (n = 6 biological replicates). **E, F)** Quantification of circuit activation in *A. thaliana* leaves with root-treated *E. coli* (left) or *P. putida* (right) at lower bacterial densities and extended incubation times (3, 5, and 7 days). Bars represent the mean of 3 independent biological replicates. Log-transformed data were analyzed by one-way ANOVA with Tukey’s HSD post hoc test. Different letters indicate statistically significant differences (p < 0.05).

Because the reporter response at OD 1 (15-fold for *E. coli* and 12-fold for *P. putida*) exceeded that of a 1 μM root-applied pC-HSL standard (2.1-fold) (**Supplementary Tables 4 and 5**), we asked whether lower bacterial densities would still produce detectable activation with longer co-culture periods. Therefore, we performed a time-course assay using bacterial suspension spanning three orders of magnitude of bacterial concentration (OD 0.01, 0.1, and 1) and quantified GFP activation after 3, 5, and 7 days. Beginning at Day 3, only the high-density inoculum (OD 1) elicited a statistically significant response, while low-density inocula remained near baseline for both *E. coli* and *P. putida*. By Days 5 and 7, the OD 1 inoculum reached maximal activation, and lower-density inocula elicited elevated signal activation with both *E. coli* and *P. putida*, particularly at OD 0.1, demonstrating that prolonged co-culture of microbes with plant roots enables robust detection even at low, soil-relevant microbial densities (**Fig. 4E and F**).

Overall, these experiments demonstrate that sentinel plants can detect and report microbially generated pC-HSL from two diverse bacterial species, establishing a functional chemical communication bridge between microbes and plants.

### Sentinel plants can detect bacterially produced pC-HSL in agricultural soil

We assessed sentinel plant performance in an agricultural soil to determine whether the platform remains functional in field-relevant conditions. We first performed a dose-response assay in agricultural soil that was mixed with sand to improve aeration and supplemented with 0 to 100 μM pC-HSL. Confocal imaging showed that the sensor response was clearly detectable in leaf tissues at 3 µM pC-HSL and increased progressively at 10, 30, and 100 µM pC-HSL (**Supplementary Fig. 12A**). Quantitative analysis of confocal microscopy images of six biological replicates revealed that the GFP intensity increased with the concentration of pC-HSL applied to the roots (**Supplementary Fig. 12B**). Sentinel plants grown in soil exhibited a 5.8- and 72-fold increase at 3 and 10 μM pC-HSL, respectively, relative to the untreated control (**Supplementary Table 6**).

We next evaluated whether bacterially produced pC-HSL could be detected in soil. Engineered bacteria were mixed with agricultural soil, and the roots of *A. thaliana* sentinel seedlings were exposed to the bacteria-inoculated soil (**Fig. 5A**). Representative confocal microscopy images of leaf tissues showed detectable GFP fluorescence in plants grown in soil inoculated with OD 1 and 10 bacteria for 72 hours (**Fig. 5B, D**). Quantitative analysis of six biological replicates showed that, relative to uninoculated controls, *E. coli* induced a 39- and 120-fold increase in reporter signal at OD 1 and 10, respectively. Similarly, *P. putida* induced a 28- and 190-fold increase at OD 1 and 10, respectively (**Fig. 5C-E, and Supplementary Table 7**). These results demonstrate that sentinel plants retain functionality in agricultural soil to report bacterially produced pC-HSL in leaves.

**Figure 5.**
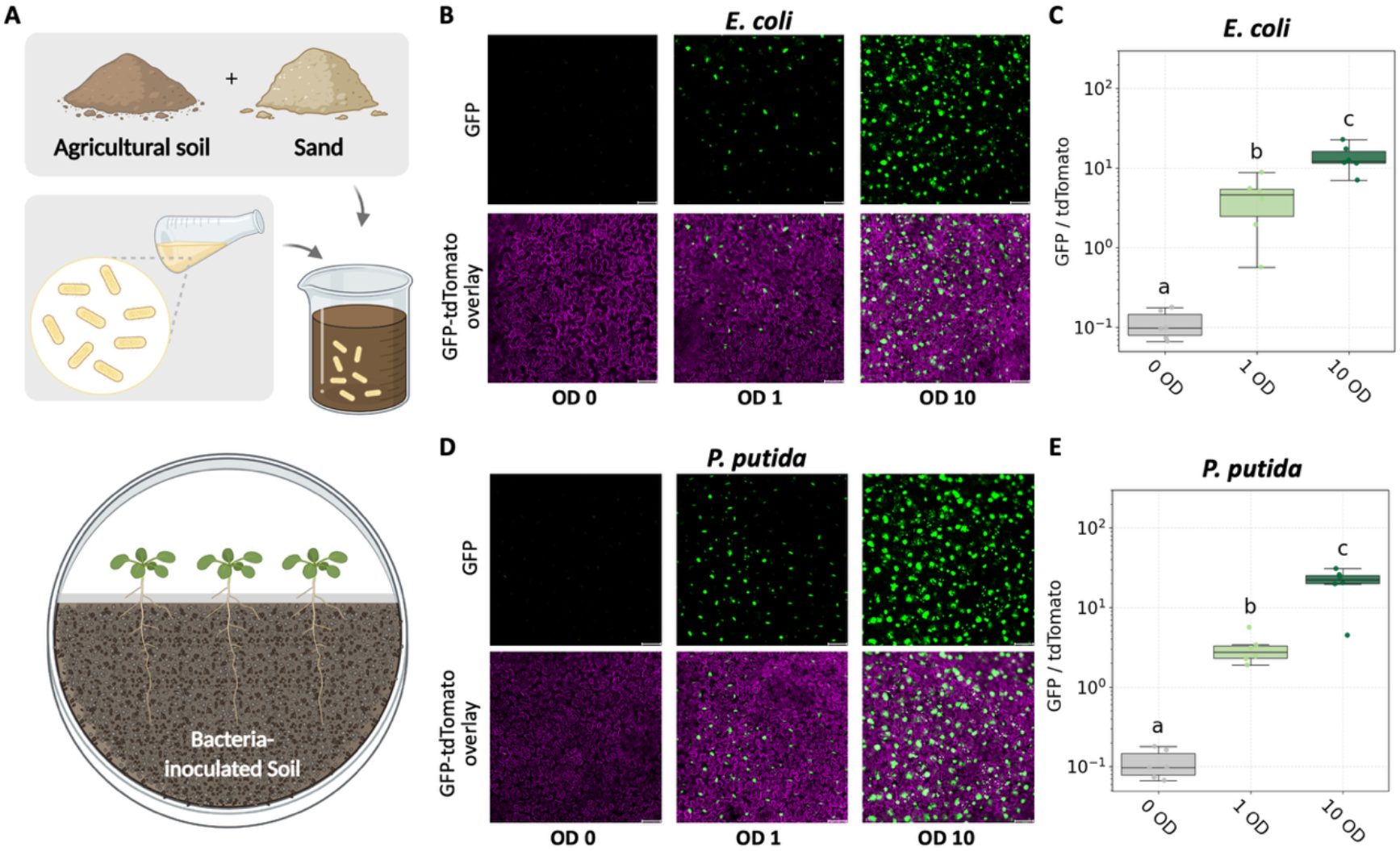
Sentinel plants detect bacterially secreted pC-HSL in soil. **A)** Schematic of the soil assay workflow. Sentinel *A. thaliana* seedlings were grown in split plates where only roots were exposed to bacteria-inoculated soil. **B, D)** Representative confocal images of GFP (green) and tdTomato (magenta) signals in *A. thaliana* sentinel plants grown in soil inoculated with engineered *E. coli* (top) or *P. putida* (bottom) suspension at OD600 ranging from 0 to 10. **C, E)** Quantification of circuit activation in *A. thaliana* leaves grown in soil inoculated with engineered *E. coli* (top) or *P. putida* (bottom) grown in soil (n = 6 biological replicates). Log-transformed data were analyzed by one-way ANOVA with Tukey’s HSD post hoc test. Different letters indicate statistically significant differences (p < 0.05).

## Discussion

In this study, we established a generalizable workflow for engineering inducible synthetic circuits in plants to provide above-ground plant signals in response to rhizosphere bacterial activity. Our findings demonstrate that plant-based biosensors can serve as effective sentinels by converting invisible soil microbial signals into visible, systemic readouts. Therefore, this work demonstrates plants’ ability to noninvasively convert rhizosphere microbial cues into visible, systemic aboveground signals, establishing plant-encoded monitoring of soil microbial activity.

We showed that transient expression screening in *N. benthamiana* using high-throughput and quantitative reporters, such as luciferase, provides a first-pass strategy to optimize circuit architecture, including operator multiplicity and transcription factor configuration. The transient assays enabled rapid identification of high-performing designs prior to committing to stable transformation. Upon transferring this optimized pC-HSL circuit into *A. thaliana* to generate stable lines, we demonstrate a dynamic range of induction in roots reaching a 2600-fold response upon treatment with 1 µM pC-HSL compared to a 36-fold response observed in transient *N. benthamiana* assays. This enhanced performance likely reflects higher expression of the RpaR-VP16 activator in the transgenic background, resulting in stronger promoter occupancy and transcriptional activation.

Alongside prior demonstrations of the functionality of canonical acyl-HSLs, our results showed that non-canonical quorum-sensing systems are also functional in plants^26^. Synthetic circuits based on quorum-sensing pathways are largely orthogonal to endogenous plant signaling pathways, making them attractive and transferable components for plant sensors across species^27^. Another quick method to validate circuit design in the species of interest is VAST (Vacuum and Sonication-Assisted Transient Expression), a transient transformation that allows versatile testing across diverse monocot and eudicot seedlings^28^.

The spatial patterns of pC-HSL-inducible reporter activation in roots indicate that longitudinal developmental gradients and radial diffusion barriers jointly shape signal perception. The endodermis forms a selective diffusion barrier by depositing suberin-rich Casparian strips that block apoplastic transport^29^. Along the longitudinal axis, stronger reporter activation near the root tip compared to more mature regions is consistent with the developmental state of the endodermis. In younger root tissues, incomplete Casparian strip formation and limited suberin deposition reduce diffusion barriers^30,31^. Along the radial axis, reporter activation at lower pC-HSL concentrations was primarily confined to epidermal and cortical layers, which are outer root tissues. In contrast, higher concentrations enabled activation in the endodermis and stele, indicating that the Casparian strip may limit signal penetration in a concentration-dependent manner. Future work can test these hypotheses using *Arabidopsis* barrier mutants.

Previous work in *Hordeum vulgare* has shown that inhibition of ATP-dependent ABC transporters by orthovanadate reduces uptake of canonical acyl-HSLs, while KCl-induced plasmodesmatal closure impairs long-distance symplastic transport, indicating that HSL movement in roots involves both active uptake and symplastic transport^26^. Further investigation will be required to determine whether similar pathways mediate pC-HSL transport in plant tissues and from root to shoots.

Beyond localized sensing within the roots, our results demonstrate that pC-HSL applied exclusively to roots can be transported systemically through the vasculature, reaching leaves via the stem and major veins. However, reporter activation was spatially heterogeneous, both among different leaves and within individual leaves. GFP fluorescence was stronger in mature leaves than in newly developing leaves and was localized in leaf parts closer to the petiole and veins. This developmental pattern likely reflects reduced accumulation of the RpaR-VP16 transcriptional activator in developing leaves, and consequently, mature leaves exhibited greater signal-activation capacity than juvenile tissues. The spatial confinement of reporter activation closer to the petiole and veins suggests that signal molecule availability in distal leaf regions may be limited.

These observations highlight an important design principle for plant-based biosensor design. Our findings indicate that sensor performance is not solely dictated by circuit architecture or signal concentration, but also by tissue developmental state and spatial competence for signal decoding. To engineer more versatile plant sensors, signal perception may be gated at the tissue or cell-type level. Such gating can involve two distinct factors: whether a signaling molecule can reach a given cell, and whether that cell can activate a transcriptional response. The former is governed by plant physiology and tissue architecture, and the latter can be genetically programmed, for example, using cell-type-specific promoters to control transcriptional regulators or essential components of the output module in target organs^32,33^. Such strategies will be essential for extending plant-based biosensing platforms to more mature plants and field-relevant conditions, where tissue heterogeneity and developmental complexity are unavoidable.

The availability of the precursor p-coumarate is a determinant of pC-HSL production by the engineered bacteria (**Supplementary Fig. 11C-D**)^25^. In soil environments, this requirement may be readily met, as p-coumarate is a phenylpropanoid derived from plant lignin and has been detected in root exudates and rhizosphere soils, where it can accumulate to levels exceeding 20 mg/kg^34,35^. Thus, plants themselves may serve as a natural source of precursor molecules. More broadly, precursor-dependent signaling systems could be leveraged to couple bacterial activity to plant presence. This strategy could be extended to enable controllable bacterial behaviors, including root-associated activation and biocontainment mechanisms.

Another consideration is the microbial density required to detect a signal. In our co-culture assays, detectable leaf-level responses were observed at an inoculation density of OD 0.1, corresponding to approximately 0.8 x 10^8^ cells mL^−1^. Given that bacterial abundances in the rhizosphere can reach up to ∼10^11^ microbial cells per gram of root, and bulk soil typically has lower densities of 10^8^-10^9^ cells per gram^36^, this observation highlights opportunities for future optimization, including enhancement of microbial signal biosynthesis, improved root colonization, and incorporation of signal amplification or relay architectures within the plant circuit to improve detection thresholds.

Lastly, we demonstrated the sentinel plant’s functionality in agricultural soil, showing responsiveness to both exogenously applied and bacterially produced pC-HSL. At OD 1 inoculation densities, reporter activation fold change in soil (38 for *E. coli* and 28 for *P. putida*) was greater than that observed in plate assays (15 for *E. coli* and 12 for *P. putida*). Despite the increased biotic and abiotic complexity of the unsterile agricultural soil matrix, engineered bacteria grew and produced pC-HSL, and the signaling molecule remained sufficiently mobile to activate the reporter circuit sensitively in the leaves of sentinel plants. Nonetheless, additional mesoscale chambers and controlled greenhouse trials will be necessary to evaluate reproducibility, stability, robustness, and long-term performance before advancing the sentinel plant platform toward field deployment.

Our findings outline the core components of a sensitive biosensing platform that reports microbial processes through a plant-encoded aboveground signal. Although this study utilized constitutive production of the signaling molecule, the bacterial module can be readily reprogrammed for inducible operation. Sensor output can be linked to endogenous bacterial transcriptional states or to promoters that respond to specific chemical or physiological cues in the rhizosphere^37^. The demonstration that aboveground tissues can decode microbial gene expression or metabolites produced exclusively in the root environment establishes an essential capability for plant-based biosensing. This root-to-shoot acropetal communication framework enables noninvasive, direct readout of rhizosphere state and microbial activity, and supports the future development of sensors that operate across plant species and soil contexts. By providing continuous, non-destructive, and scalable access to key microbial dynamics, this sentinel-plant approach can advance long-term soil management, precision agriculture, and ecosystem monitoring.

## Methods

### Construct design and cloning

The core pC-HSL-inducible cassette was kindly provided by the Christopher Voigt lab^16^. Plasmids were assembled with the Loop Assembly method using BsaI- and SapI-mediated Golden Gate reactions (NEB catalog numbers R0569S and E1601L, respectively) and cloned into pCAMBIA backbones. All plasmids were propagated in NEB Turbo *Escherichia coli* competent cells (NEB catalog no. C2984I) and sequence-verified by nanopore sequencing (Plasmidsaurus). Plasmids generated in this work are listed in **Supplementary Table 8**.

### Plant growth conditions

*A. thaliana* (Col-0) seeds were surface-sterilized by exposure to chlorine gas. Chlorine gas was produced by mixing 100 mL of bleach and 3 mL of 6N hydrochloric acid. The mixture was placed in a desiccator box within a chemical fume hood, and the seeds were exposed to chlorine gas for at least 3 hours. Seeds were germinated on ½ MS plates containing 0.5x Murashige and Skoog Basal Salt Mixture (M524, PhytoTech Labs), 0.05% MES (w/v), 0.7% Phytagel (P8169, Millipore Sigma), pH 5.7, and incubated in a growth chamber at 22°C with a 16/8 h light/dark cycle. *Nicotiana benthamiana* plants were grown in controlled growth chambers (Conviron) under long-day conditions (16 h light / 8 h dark) set to 22 °C.

### N. benthamiana transient luciferase assay

*Agrobacterium tumefaciens* GV3101 strains harboring circuit constructs were cultured overnight at 30 °C with shaking at 200 rpm in 2×YT medium (ThermoFisher 22712020) supplemented with 10 µg mL^−1^ rifampicin (Sigma-Aldrich R3501), 20 µg mL^−1^ gentamicin (Sigma-Aldrich G1264), 50 µg mL^−1^ tetracycline (Sigma-Aldrich T7660), and 50 µg mL^−1^ kanamycin (Sigma-Aldrich K1637). Cultures were pelleted by centrifugation at 4,000 g for 5 min and resuspended in infiltration buffer (10 mM MES, pH 5.7; 10 mM MgCl_2_; 200 µM acetosyringone) to a final OD_600_ of 0.4. Resuspended cultures were incubated at room temperature with shaking at 120 rpm for 4 h to induce virulence gene expression prior to infiltration.

Agrobacterium suspensions were infiltrated into the abaxial surface of fully expanded leaves of 3-4-week-old *N. benthamiana* plants using a needleless syringe. At 48 h post-infiltration, the same leaf areas were infiltrated with pC-HSL (Millipore Sigma 07077-50MG) at concentrations ranging from 0 to 10 µM. Leaf tissue was harvested 48 h after pC-HSL treatment for luciferase analysis.

For luminescence measurements, three leaf discs were excised from infiltrated leaf areas using a 1/4″ radius leaf punch and placed into 2 mL microcentrifuge tubes (Fisher Scientific 15-340-162) containing 2.8 mm ceramic beads (Fisher Scientific 15-340-160). Samples were immediately flash-frozen in liquid nitrogen and homogenized using a mini bead-beater tissue homogenizer (Fisher Scientific 15-340-164) for two 20-s cycles at maximum speed. Following homogenization, 200 µL of 1× Cell Lysis Buffer (Promega E1531) was added to each tube, and samples were vortexed vigorously until fully resuspended. Lysates were incubated on ice for 10 min and clarified by centrifugation at 12,000 g for 5 min to pellet cellular debris.

Luciferase activity was measured using the Nano-Glo® Luciferase Assay System (Promega N1110) according to the manufacturer’s instructions with minor modifications. Briefly, 30 µL of clarified lysate was mixed with 60 µL of Nano-Glo reagent in a 96-well plate (Fisher Scientific 07-200-627). Luminescence was recorded using a Tecan Spark plate reader at 25 °C with a 1 s integration time per well. GeNL and NanoLuc signals were collected using 520–590 nm and 430–485 nm emission filters, respectively. Signal deconvolution coefficients were calculated by measuring total, GeNL-only, and NanoLuc-only luminescence and applying the Promega ChromaLuc Technology calculator (Technical Manual TM062).

### A. thaliana floral spray transformation and selection

*Agrobacterium tumefaciens* cultures (50 mL) harboring the appropriate plant expression constructs were grown to the stationary phase, pelleted by centrifugation at 4,000 g for 10 min, and resuspended in 5% (w/v) sucrose supplemented with 0.02% (v/v) Silwet L-77 (Fisher Scientific NC1765106). Flowering *A. thaliana* plants (6-7 weeks old) were uniformly sprayed with the *Agrobacterium* suspension until runoff^38^. Plants were covered overnight with a light-blocking dome to maintain humidity and minimize light exposure, then returned to standard growth conditions.

Upon seed maturation, T1 seeds were screened for FAST-Red fluorescence, and positive seeds were picked out and germinated on ½ MS agar plates supplemented with 5 µg mL^−1^ glufosinate ammonium (BASTA; Millipore Sigma 45520-100MG) to select for resistant transformants. Multiple independent lines were screened for reporter responsiveness, and a single high-performing homozygous line was selected for downstream experiments.

### Confocal microscopy and fluorescence quantification

Plant samples were mounted in distilled water and imaged immediately using a Leica Stellaris 8 confocal laser-scanning microscope equipped with an HC PL APO 10×/0.40 CS2 objective, housed in the Biological Imaging Facility (BIF) at Caltech. A pulsed white-light laser at 488 nm was used to excite GFP, and emission spectra were collected between 500 and 540 nm. The pulsed white light laser set to 561 nm was used to excite tdTomato fluorescence, and emission was collected between 580 and 630 nm. For single-frame imaging, one optical z-section was acquired per field of view with the pinhole set to 10 AU. For z-stack acquisition, optical sections were collected sequentially with the pinhole also set to 1 AU and a z-step size of 5 µm. Identical laser power, detector gain, and offset settings were maintained for all samples within each experiment. Images were exported from Leica LAS X software (v4.3) as 16-bit TIFF files.

Each data point represents the average across multiple FOVs from a single biological replicate corresponding to a single plant. 6 to 9 biological replicates were used in these experiments (see Figure captions). For *N. benthamiana* leaves, a leaf punch adjacent to the infiltration site was collected. The mean fluorescence intensity of each biological replicate is calculated from 4 random non-overlapping FOVs within the same punch. For *A. thaliana* leaves, the mean fluorescence intensity of each biological replicate is calculated from 3 random non-overlapping FOVs of the same leaf (**Supplementary Figure 9)**. For *A. thaliana* roots, each datapoint corresponds to a single FOV. The fluorescence intensity of each FOV is calculated in Fiji/ImageJ using the “integrated density” option, which measures the sum of pixel intensity values within a selected region of interest, with the region set to the entire FOV. Fold-change was calculated as the ratio of integrated GFP to tdTomato fluorescence intensity, multiplied by 100.

### Chemical induction of sentinel plant roots with pC-HSL

Transgenic *A. thaliana* seedlings were grown vertically on ½ MS agar plates for 10 days. Seedlings were then transferred to ½ MS agar plates supplemented with defined concentrations of pC-HSL. Plates were maintained in a vertical orientation to preserve root architecture. After 48 h of induction, seedling roots were imaged by confocal microscopy as described.

### Chemical induction of sentinel plant leaves with pC-HSL

Transgenic *A. thaliana* seedlings were grown vertically on ½ MS agar plates for 10 days. Seedlings were then transferred to vertically oriented split agar plates (I-plates; VWRU25384-344). The lower compartment of the petri dish contained ½ MS agar supplemented with defined concentrations of pC-HSL, while the upper compartment contained ½ MS agar without any inducer. Roots were positioned in the lower compartment, and shoots were exclusively placed in the upper compartment, ensuring that leaves did not contact pC-HSL-containing media. Plates were maintained in a horizontal orientation. After 72 h of induction, seedling leaves were imaged by confocal microscopy.

### Construction of pC-HSL-producing E. coli and Pseudomonas putida strains

To construct the pC-HSL production cassette, the three-gene pC-HSL production operon from Du et al. ^19^ was commercially synthesized as a dsDNA fragment with BsaI overhangs following standard CIDAR MoClo Assembly format^39^. This operon was then combined with the J23101 constitutive promoter and a transcriptional terminator and cloned into a kanamycin-resistant p15a vector using 3G assembly^40^. The resulting plasmid was then transformed into chemically competent *E. coli* JM109 cells by adding 1 µL of plasmid assembly product to 50 µL of competent cells and incubating on ice for 10 minutes, after which a 42C heat shock was applied for 1 minute. After 3 minutes of additional incubation on ice, 1 mL of SOC media was added to the cells, and they were outgrown for 1 hour at 37C shaking at 250 rpm before plating on LB agar plates with 50 µg/mL kanamycin selection. Colonies from these plates were then grown up and miniprepped to isolate the plasmid, which was validated using whole-plasmid sequencing.

The *E. coli* pC-HSL production strain was constructed by transforming the resulting plasmid into chemically competent *E. coli* BW25113 cells using the heat shock protocol described above. Similarly, the *P. putida* pC-HSL production strain was constructed in the following way. Electrocompetent *Pseudomonas putida* KT2440 cells were thawed on ice immediately prior to transformation. Purified plasmid DNA (100-500 ng) was added to 40 µL of competent cells, and the mixture was gently mixed. The cell and plasmid mixture was transferred to a pre-chilled 0.1 cm gap electroporation cuvette (Bio-Rad 652089) and incubated on ice for 10 min. Electroporation was performed using a Bio-Rad MicroPulser electroporator. Immediately following the pulse, 500 µL of SOC medium was added to the cuvette, and the cells were transferred to a 1.5 mL microcentrifuge tube. Cells were recovered by incubation at 30 °C with shaking at 200 rpm for 2 h, then plated on selective agar and incubated overnight at 30 °C.

### Construction of pC-HSL-responsive E. coli biosensor

We constructed a pC-HSL-responsive GFP biosensor by using 3G Assembly on parts sourced from the CIDAR MoClo Part Extension Kit Volume 1^41^. Notably, the specific sequences of the pC-HSL-responsive transcription factor RpaR and its cognate promoter P_rpa_ were from previously reported engineered variants^42^. The resulting plasmid assembly was transformed into *E. coli* JM109 cells and validated as described above. These strains were used directly as the pC-HSL biosensor.

### Measurement of pC-HSL production from engineered E. coli and P. putida

The *E. coli* pC-HSL sensor strain was grown overnight in M9CA medium (Teknova M8010) at 30 °C with shaking at 200 rpm. On the day of the bioassay, the overnight culture was diluted 1:100 into 5 mL of M9CA medium and incubated for 3 h at 30 °C, shaking at 200 rpm to reach the exponential phase. This culture was then diluted an additional 1:100, and 200 µL aliquots were dispensed into a 96-well plate (Fisher Scientific FB012931) for downstream assays.

Engineered *E. coli and P. putida* strains were revived from glycerol stocks and grown overnight in M9CA medium at 30 °C with shaking at 200 rpm. Cultures were then diluted to a starting OD_600_ of 0.01 in 3 mL of M9CA medium, with or without 100 µM p-coumarate (Millipore Sigma C9008), in a 24-deep-well plate (ThermoFisher Scientific 95040470). Cultures were incubated at 30 °C with shaking at 200 rpm, and 100 µL samples were collected from the same well every 12 h. Samples were centrifuged to pellet bacteria, and the clarified supernatants were retained at -20 °C for later analysis.

On the day before the bioassay for measuring pC-HSL concentrations from these samples, the *E. coli* pC-HSL sensor strain was revived from glycerol stocks in grown overnight in M9CA medium at 30 °C with shaking at 200 rpm. This overnight culture was then diluted 1:100 into 5 mL of M9CA medium and incubated for 3h at 30 °C and 200 rpm shaking to reach the exponential growth phase. This culture was then diluted an additional 1:100 into 200 µL aliquots in a 96 well plate (Fisher Scientific FB012931).

Purified pC-HSL (Sigma-Aldrich 07077) was used to generate a 12-point standard curve spanning 1 nM to 100 µM pC-HSL which was applied to three technical replicates of the *E. coli* biosensor to calibrate its response to known concentrations of pC-HSL. In the same plate, 2 µL of each collected supernatant sample was added to the prepared sensor strain solution. sfGFP fluorescence was measured every 20 min for 24 h of incubation at 30 °C with shaking using a Tecan Spark plate reader. Fluorescence values from the supernatants from the production strains were compared against the response to the calibration curve via linear regression to calculate the observed pC-HSL in the sample wells, which is 1/100 of the concentration in the original supernatant. In cases where fluorescence values from the supernatants fell outside of the linear range of the biosensor response, the experiments were repeated with the supernatant samples undergoing further dilution as needed to have the biosensor response lie in the linear range of the calibration curve.

### Plant-microbe co-culture assay

Engineered bacterial strains were grown overnight in 5 mL M9CA medium supplemented with p-coumarate at 30 °C with shaking. The following day, 200 µL of the overnight culture was inoculated into 20 mL of fresh M9CA medium supplemented with 100 µM of p-coumarate and grown for an additional 24 h. Bacterial cultures were harvested by centrifugation at 4,000 g for 5 min and resuspended in fresh M9CA medium to the desired optical density.

Transgenic *A. thaliana* seedlings were grown vertically on ½ MS agar plates for 10 days. Seedlings were then transferred to vertically oriented split agar plates. The lower compartment of the petri dish contained ½ MS agar supplemented with 100 µM of p-coumarate. A total volume of 400 µL of the bacterial suspension was applied evenly to the lower compartment. Roots were placed in the lower compartment, and shoots were placed exclusively in the upper compartment, ensuring that leaves did not contact the bacterial solution. Plates were maintained in a horizontal orientation. After 72 h of induction, seedling leaves were imaged by confocal microscopy.

### Soil preparation

Agricultural soil was collected from a post-harvest field site at the University of California Riverside Agriculture Experiment Station. Soil was collected and sieved through a 1mm sieve and stored at 4C until the start of the experiment. For plant experiments, the soil mixture consisted of 10% (v/v) agricultural soil and 90% (v/v) riverbank sand (Amazon). Water was added at 10% (v/w) relative to total soil mass to achieve optimal moisture content for plant growth.

### Chemical induction of sentinel plant leaves with pC-HSL in soil

Transgenic *A. thaliana* seedlings were grown vertically on ½ MS agar plates for 10 days. Seedlings were then transferred to a split petri dish. The lower compartment of the petri dish contained 20 g of soil supplemented with defined concentrations of pC-HSL, while the upper compartment contained 20 g of soil without any inducer. Roots were positioned in the lower compartment, and shoots were exclusively placed in the upper compartment. Plates were maintained in a horizontal orientation. After 72 h of induction, seedling leaves were imaged by confocal microscopy.

### Plant-microbe co-culture assay in soil

The co-culture medium consisted of M9CA medium and MS medium mixed at a 1:1 (v/v) ratio and supplemented with 100 μM p-coumarate. Engineered bacterial strains were grown overnight in 10 mL co-culture medium at 30 °C with shaking. The following day, 4 mL of the overnight culture was inoculated into 20 mL of co-culture medium and grown for approximately 6 h to reach the exponential phase. Bacterial cultures were harvested by centrifugation at 4,000 g for 5 min and resuspended in fresh co-culture medium to the desired optical density.

Transgenic *A. thaliana* seedlings were grown vertically on ½ MS agar plates for 10 days. Seedlings were then transferred to a vertically oriented split petri dish. The lower compartment of the petri dish contained 20 g of soil watered with a bacterial solution, while the upper compartment contained 20 g of soil without any bacterial solution. Roots were placed in the lower compartment, and shoots were placed exclusively in the upper compartment, ensuring that leaves did not contact the bacterial solution. Plates were maintained in a horizontal orientation. After 72 h of induction, seedling leaves were imaged by confocal microscopy.

## Data Availability Statement

All data are included either in the manuscript or supplementary files.

## Author Contributions

Conceptualization: YW, JPM, GSD

Methodology: YW, JPM, PT

Data Acquisition: YW, JPM, EL, PT

Supervision: GSD

Writing: YW, GSD

Funding Acquisition: JPM, EM, GSD

## Acknowledgements

The authors thank the RSI Impact Grant team, Bruce Hay, Niles Pierce, Shuwen (Eric) Lei, and Mikhail Hanewich-Hollatz for valuable discussions and feedback. The authors acknowledge the Caltech Biological Imaging Facility, support from the Caltech Beckman Institute, and the Arnold and Mabel Beckman Foundation for providing access to the Leica STELLARIS 8 inverted confocal microscope. Schematics were created with BioRender.com.

