## Supplementary Figures for "Sentinel Plants Enable Aboveground Detection of Belowground Soil Microbial Activity"

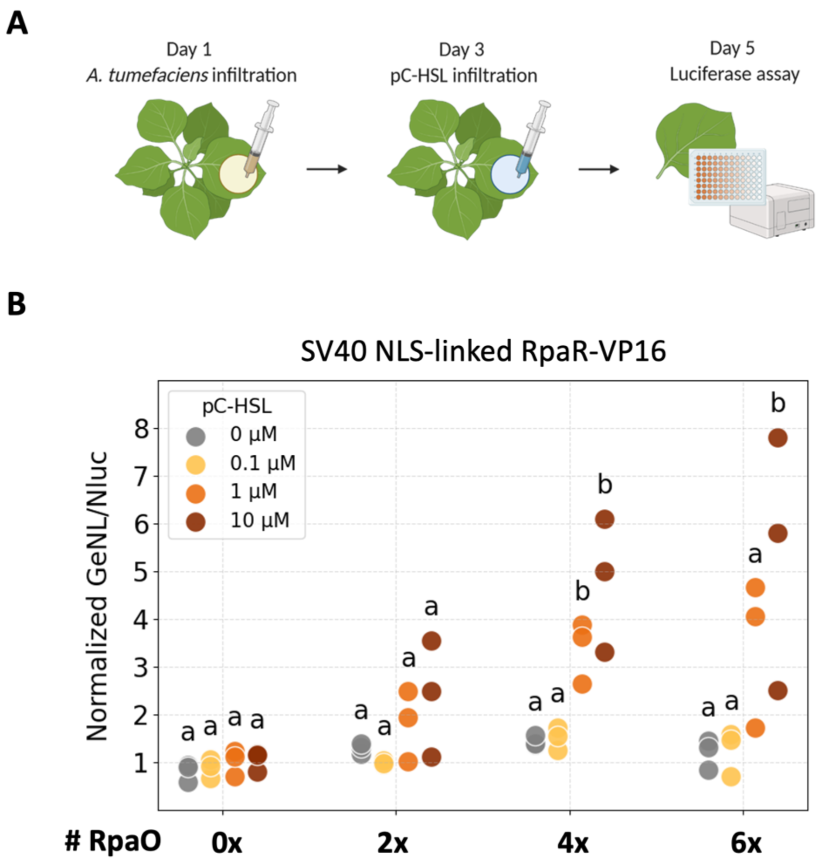


**Supplementary Figure 1. Ratiometric luciferase readout of pC-HSL-inducible circuit activity in *Nicotiana benthamiana*. A)** *N. benthamiana* leaves were first infiltrated with *Agrobacterium tumefaciens* carrying the reporter circuit, followed by infiltration of pC-HSL on Day 3. Luciferase signal is quantified on Day 5 with a plate reader. **B)** Normalized GeNL/NanoLuc activity for a library of circuit variants containing the SV40 NLS-linked RpaR-VP16 treated with 0, 0.1, 1, or 10 µM pC-HSL. Three independent biological replicates are shown. Log-transformed data were analyzed by one-way ANOVA with Tukey's HSD post hoc test. Different letters indicate statistically significant differences ($p < 0.05$).

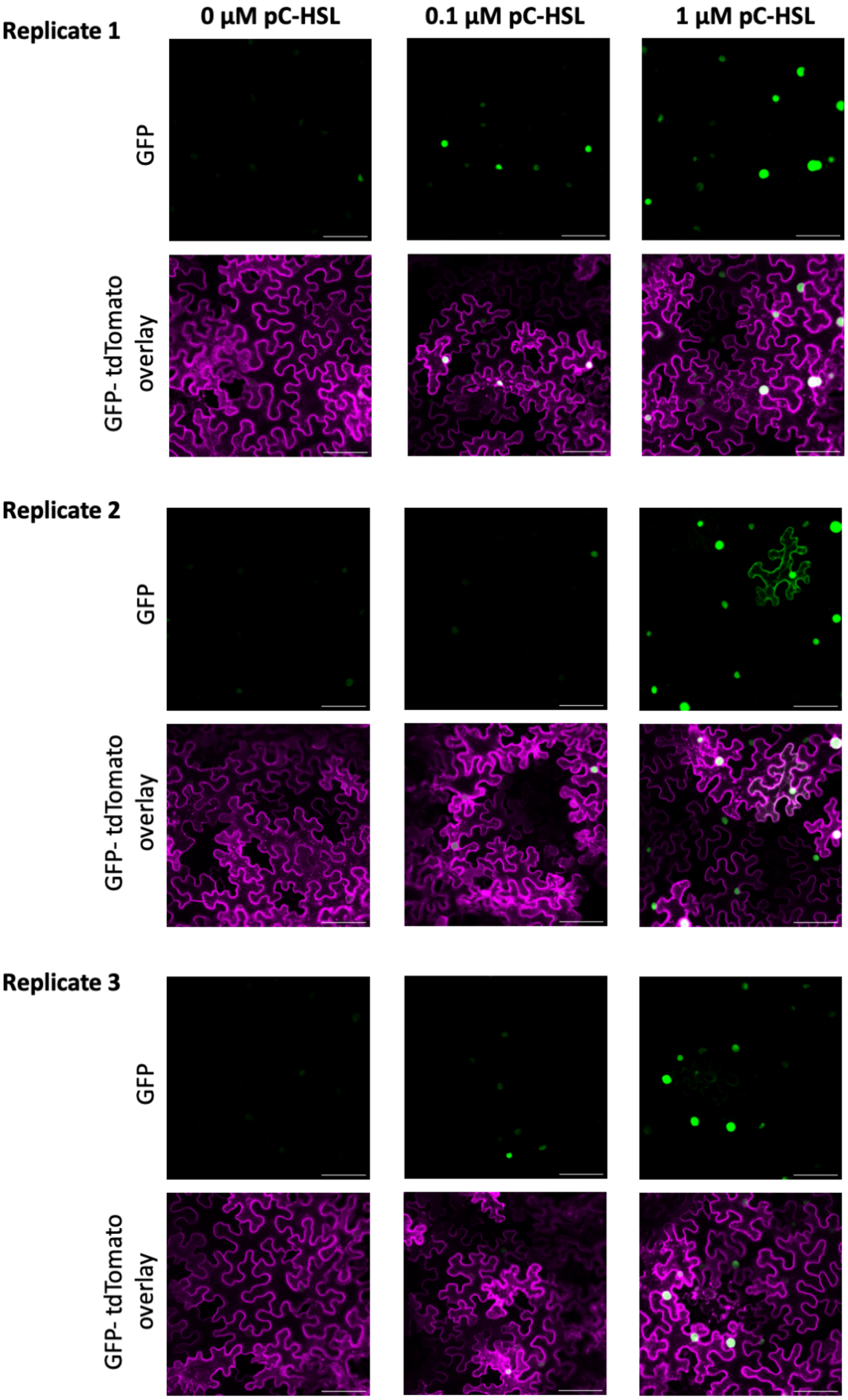


**Supplementary Figure 2. Replicate confocal microscopy images of pC-HSL-induced GFP activation in *N. benthamiana.*** Representative confocal microscopy images from three independent biological replicates showing GFP (top) and GFP/tdTomato overlay (bottom) in *N. benthamiana* leaves transiently expressing the optimized pC-HSL-inducible circuit and treated with 0, 0.1, or 1 µM pC-HSL. All scale bars, 100 µm.

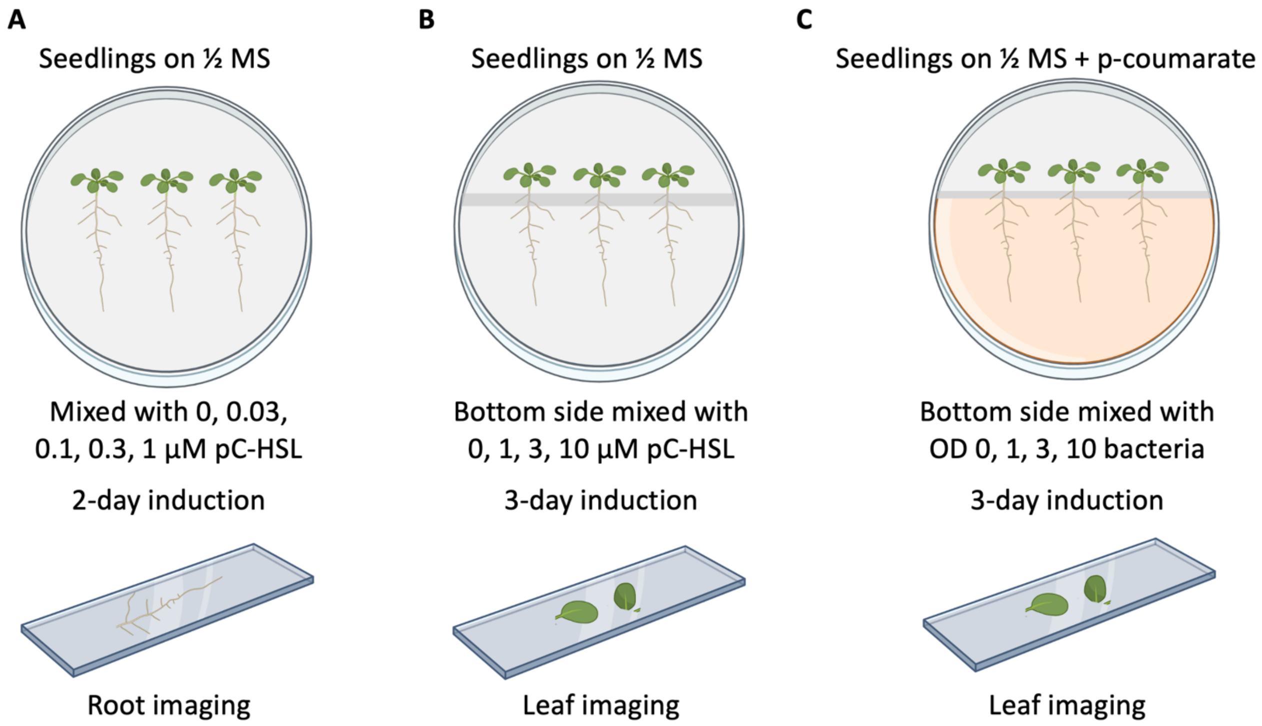


**Supplementary Figure 3. Experimental setup for pC-HSL induction assays. A)** Root induction assay. 10-day-old seedlings were transferred to ½ MS agar plates supplemented with pC-HSL (0, 0.03, 0.1, 0.3, or 1 µM). Roots were imaged by confocal microscopy after 2 days. **B)** Leaf induction assay. 10-day-old seedlings were grown on vertically oriented split ½ MS agar plates, with pC-HSL (0, 1, 3, or 10 µM) applied exclusively to the lower compartment containing the roots. Leaves were physically separated and imaged by confocal microscopy after 3 days. **C)** Root-microbe co-culture assay. Ten-day-old seedlings were grown on split ½ MS agar plates supplemented with p-coumarate, and engineered bacterial suspensions (OD600 = 0, 1, 3, or 10) were applied only to the lower compartment. Leaves were imaged by confocal microscopy after 3 days.

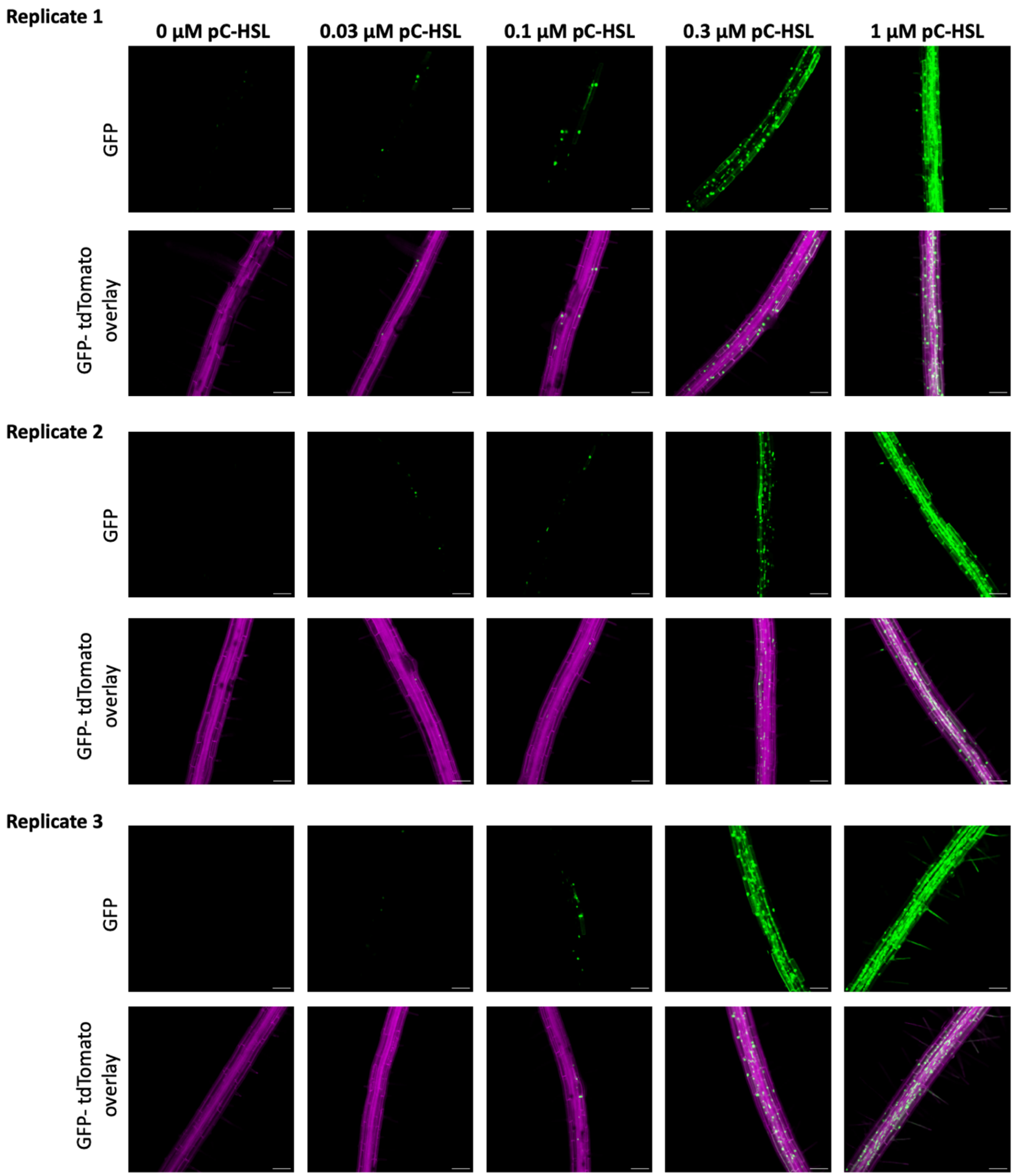


**Supplementary Figure 4. Replicate representative confocal images of pC-HSL-induced GFP activation in roots of stably transformed *A. thaliana* line #1.** Confocal microscopy images from three independent biological replicates showing GFP (top) and GFP/tdTomato overlay (bottom) in roots of stably transformed *A. thaliana* seedlings treated with 0, 0.03, 0.1, 0.3, or 1 µM pC-HSL. Scale bars, 100 µm.

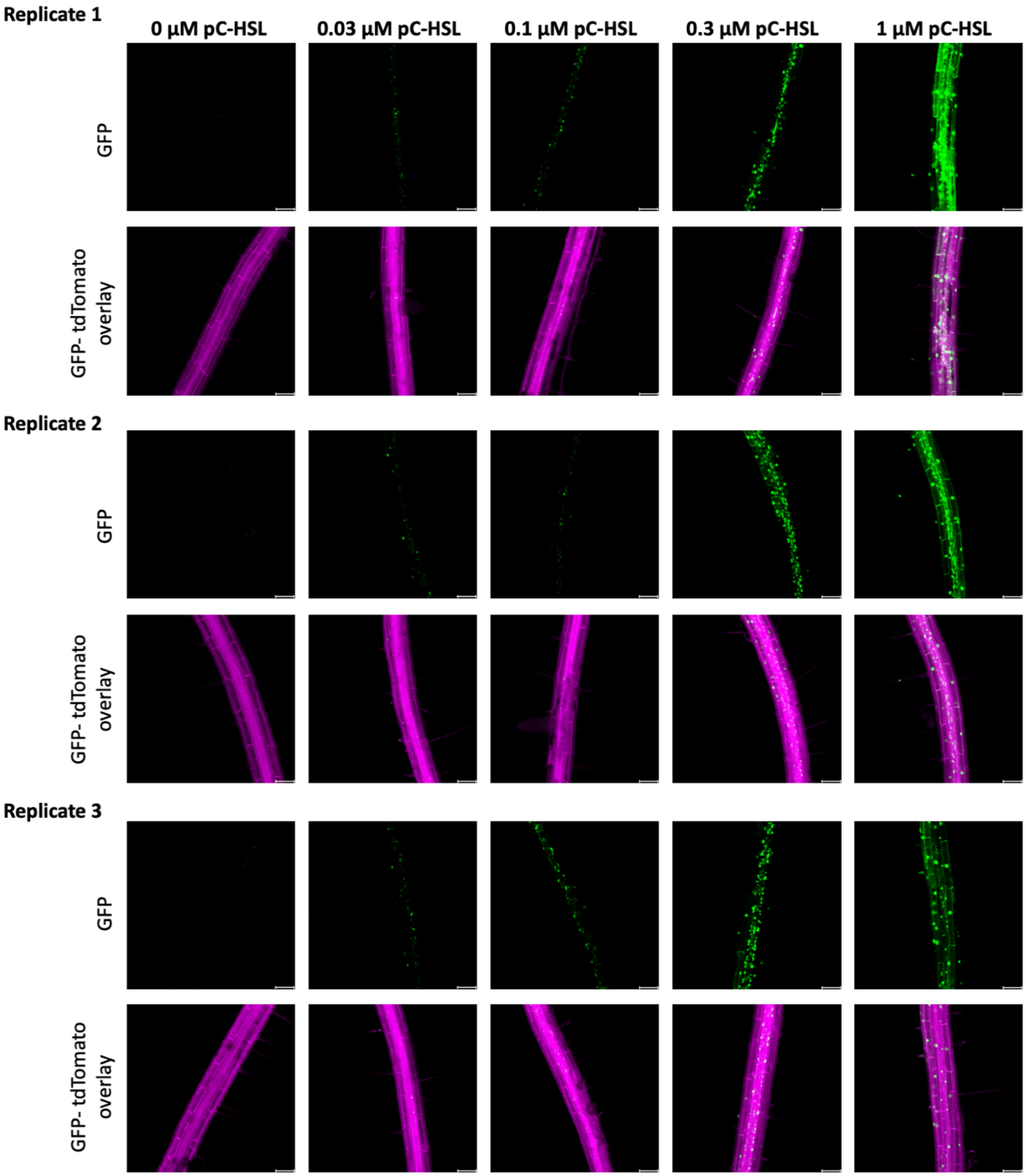


**Supplementary Figure 5. Replicate representative confocal images of pC-HSL-induced GFP activation in roots of stably transformed *A. thaliana* line #2.** Confocal microscopy images from three independent biological replicates showing GFP (top) and GFP/tdTomato overlay (bottom) in roots of stably transformed *A. thaliana* seedlings treated with 0, 0.03, 0.1, 0.3, or 1 µM pC-HSL. Scale bars, 100 µm.

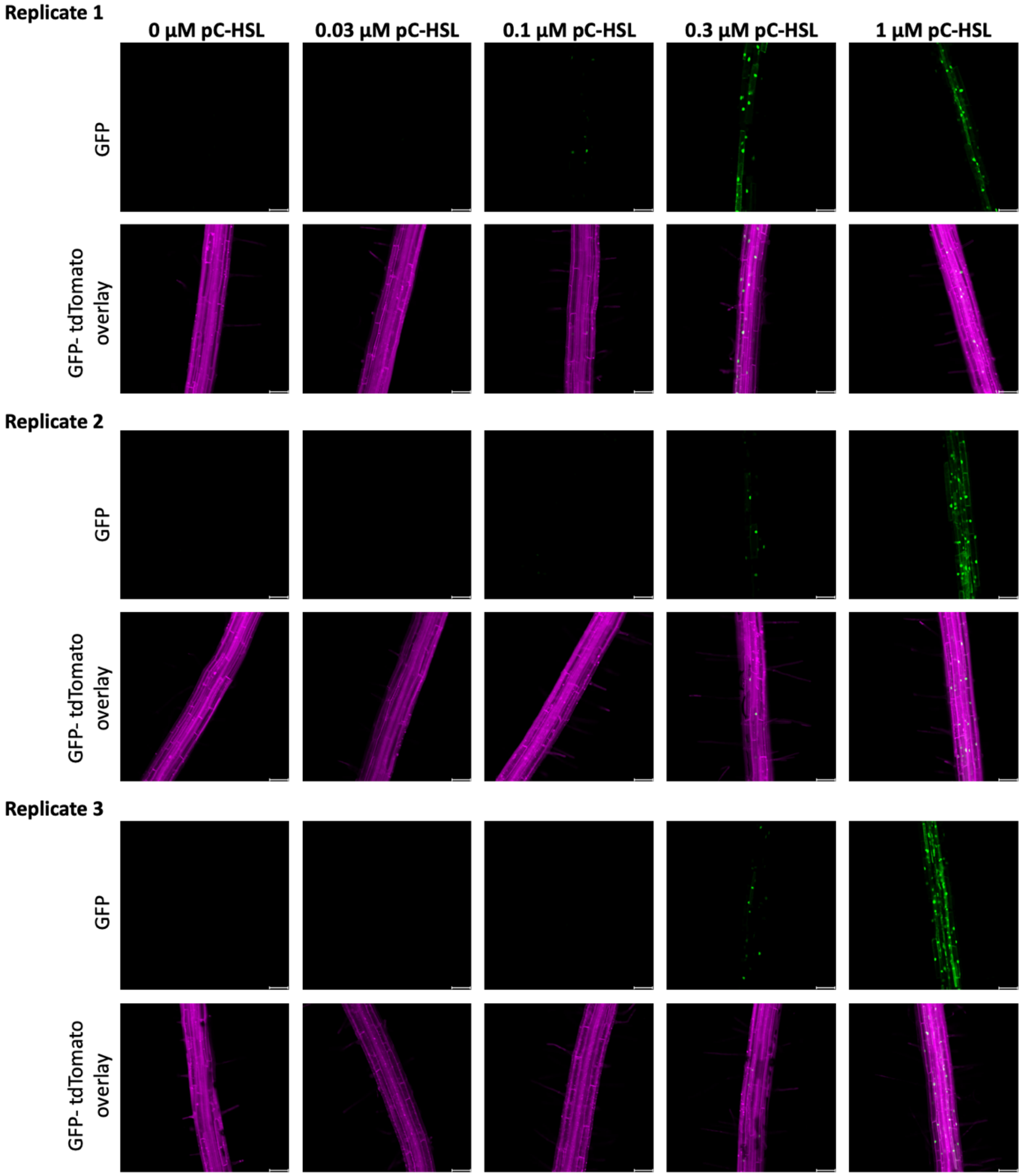


**Supplementary Figure 6. Replicate representative confocal images of pC-HSL-induced GFP activation in roots of stably transformed *A. thaliana* line #3.** Confocal microscopy images from three independent biological replicates showing GFP (top) and GFP/tdTomato overlay (bottom) in roots of stably transformed *A. thaliana* seedlings treated with 0, 0.03, 0.1, 0.3, or 1 µM pC-HSL. Scale bars, 100 µm.

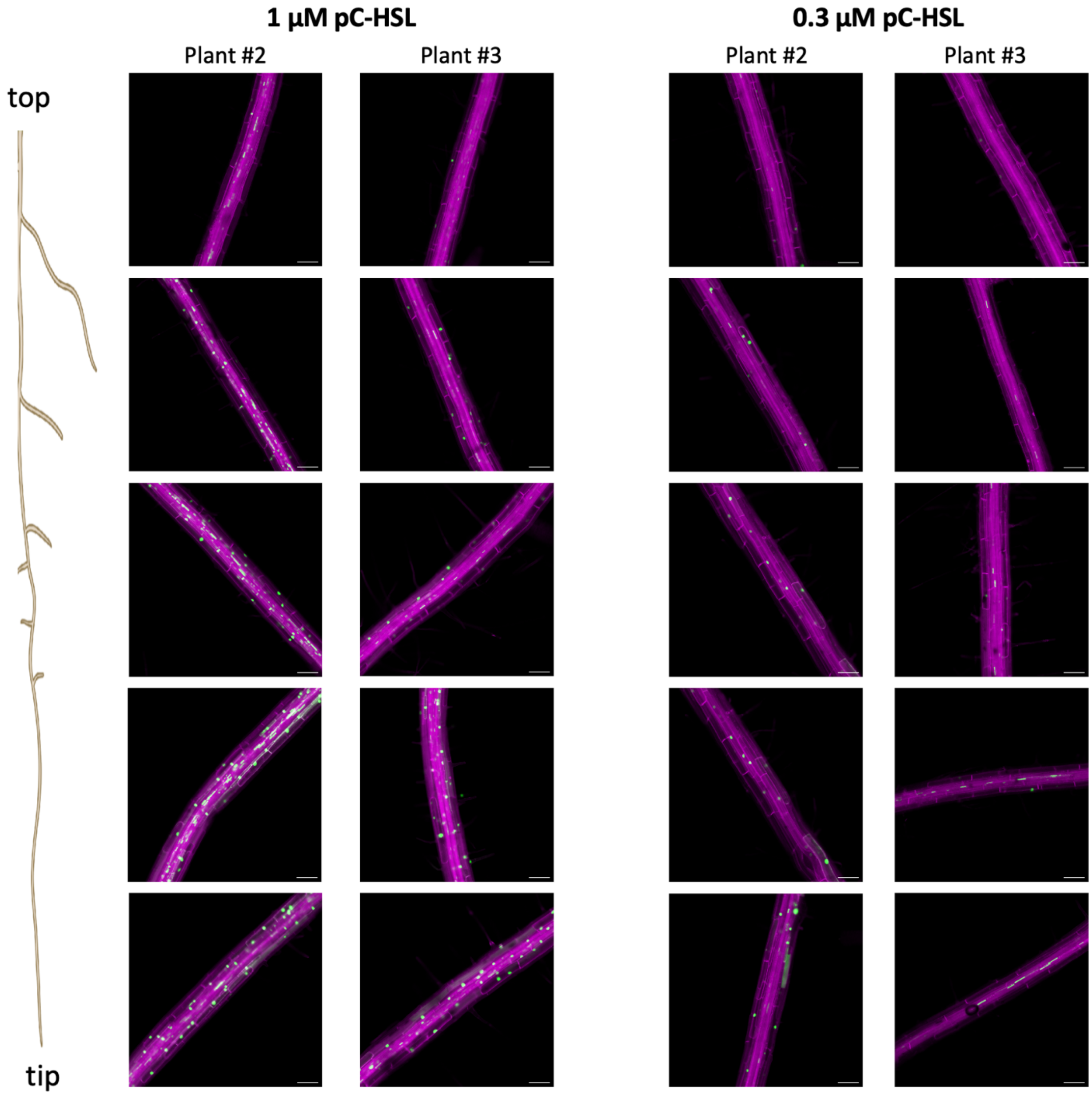


**Supplementary Figure 7. Spatial profile of pC-HSL-induced circuit activation along the root longitudinal axis.** Additional images of the spatial profile of GFP activation along the primary root treated with 1 µM and 0.3 µM pC-HSL in two other independent plants. Images are shown from bottom to top as fields of view progressing from the root tip toward the elongation and maturation zones. Scale bars, 100 µm.

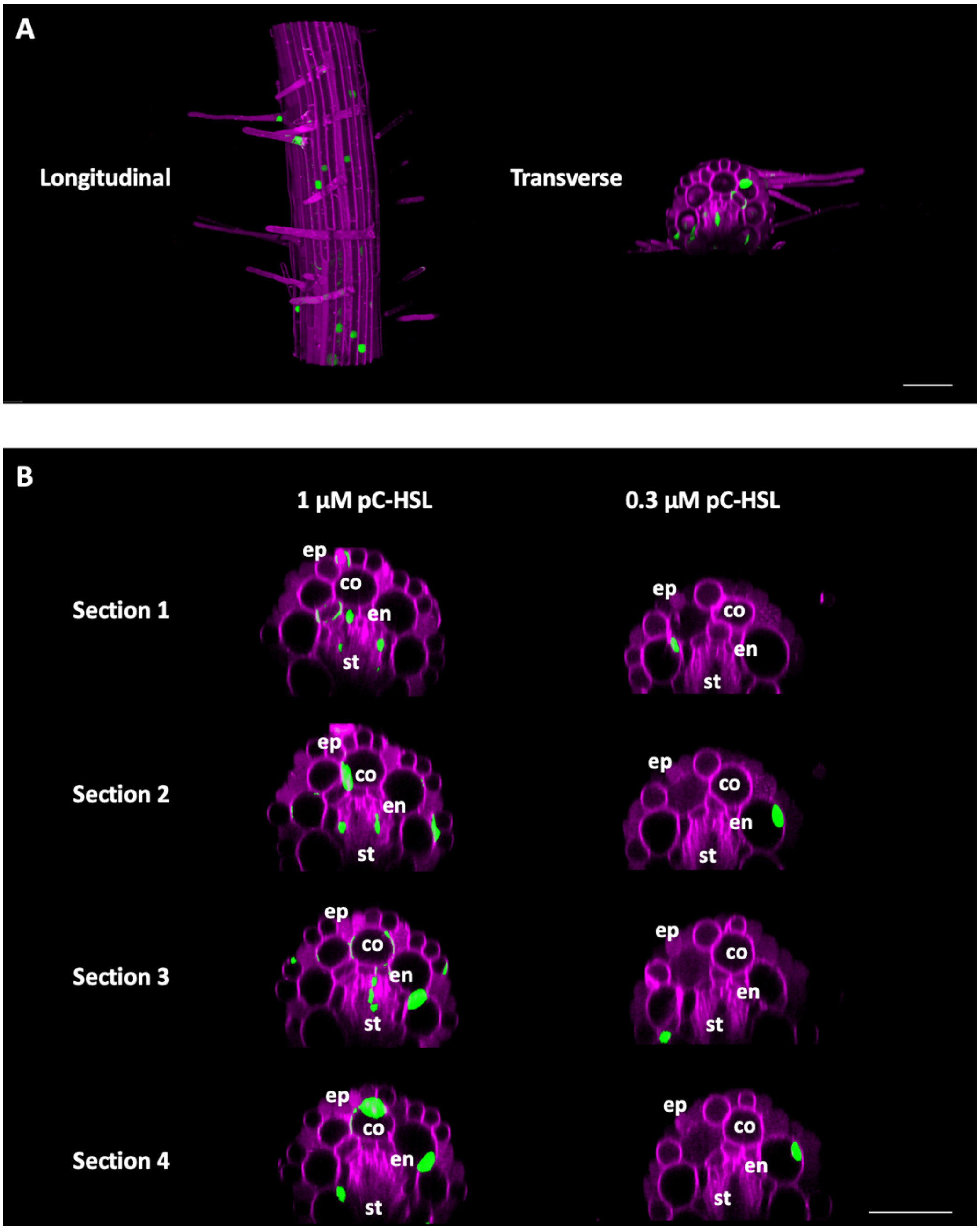


**Supplementary Figure 8. Spatial profile of pC-HSL-induced circuit activation along the root radial axis.** Representative images of simulated transverse optical sections showing the distribution of circuit activation across cell layers in roots treated with 1 μM and 0.3 μM pC-HSL. Sections represent different parts of the root. ep: epidermis, co: cortex, en: endodermis, st: stele. Scale bars, 100 μm.

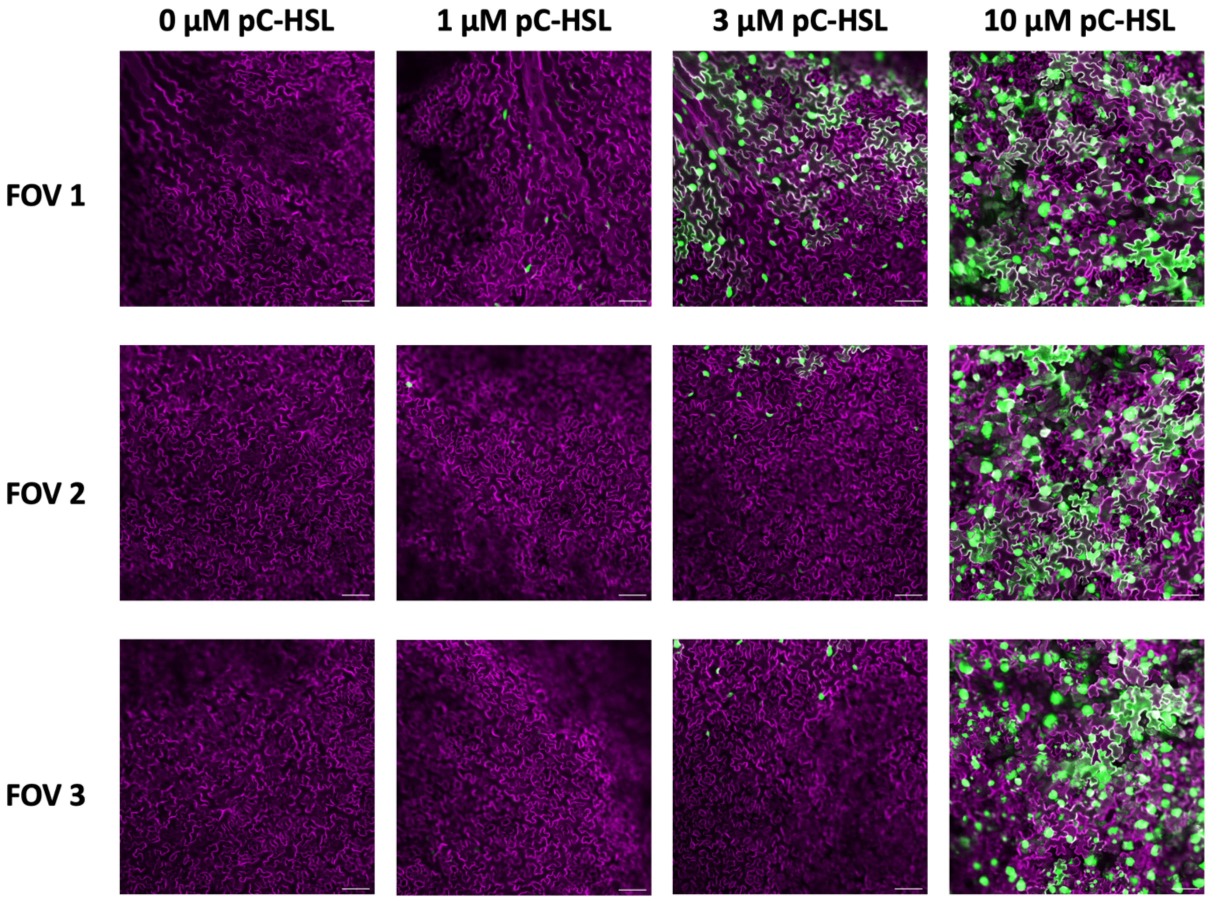


**Supplementary Figure 9. Representative fields of view used for quantifying leaf GFP activation in stably transformed *A. thaliana* (Line #1).** Confocal images from three representative fields of view (FOV) showing GFP (green) and tdTomato (magenta) fluorescence in *A. thaliana* leaves with root-treated pC-HSL at 0, 1, 3, or 10 μM. FOV1 is close to the vasculature, and FOV2 and FOV3 are in non-vascular regions. Signals from the three fields of view were averaged to quantify systemic reporter activation. Scale bars, 100 μm.

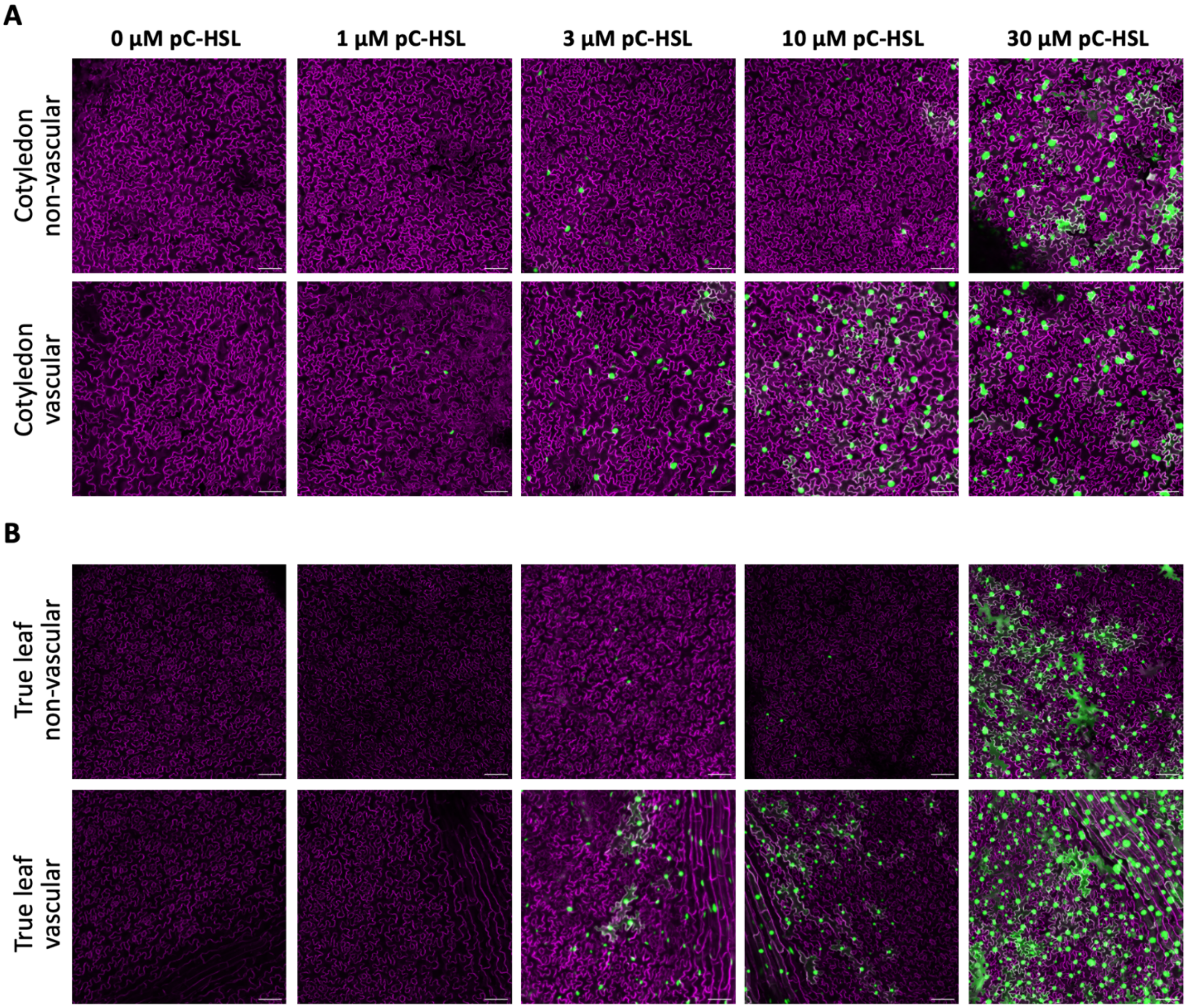


**Supplementary Figure 10. Sensor response comparison in vascular and non-vascular areas of cotyledon and young true leaves of root-treated sentinel plants.** Fluorescence signal activation differs across leaf developmental stages, with cotyledons showing higher GFP levels than newly emerging leaves with root-treated pC-HSL at concentrations ranging from 0 to 30 µM. Scale bars, 100 µm.

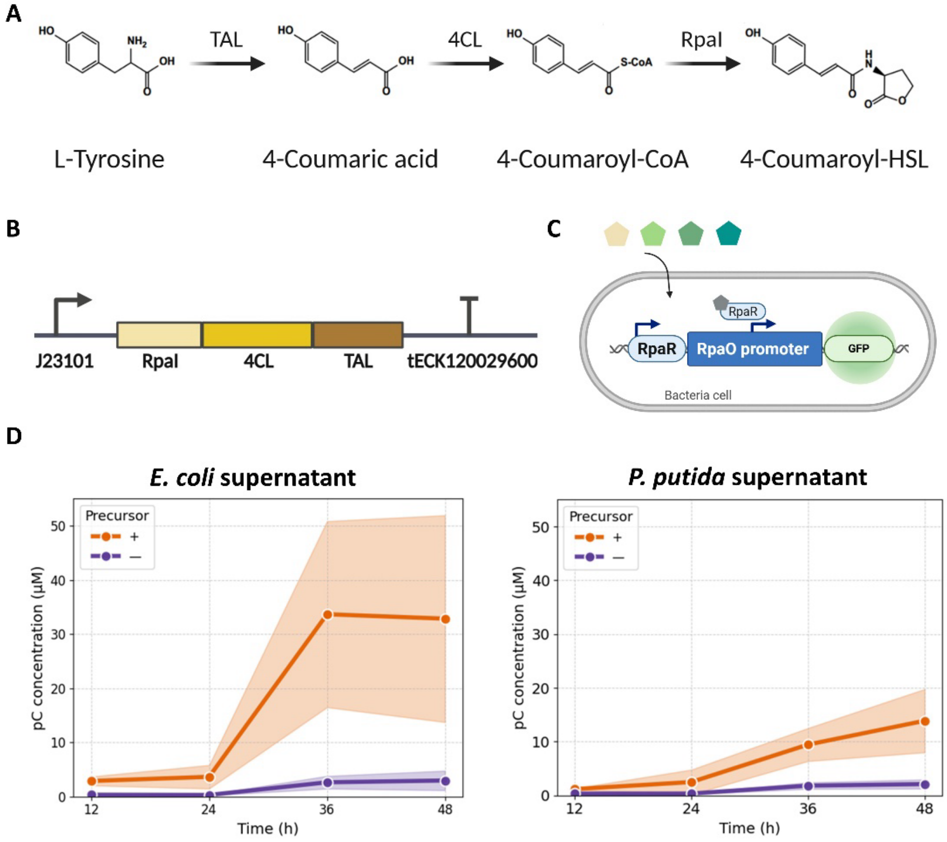


**Supplementary Figure 11. Synthetic pathway and reporter system for microbial production and detection of pC-HSL. A)** Construct map of the synthetic operon used for pC-HSL biosynthesis in bacteria. The operon contains the constitutive promoter J23101 driving expression of RpaI (p-coumaroyl-homoserine lactone synthase), 4CL (4-coumarate-CoA ligase), and TAL (tyrosine ammonia lyase), followed by the strong intrinsic terminator tECK120029600. **B)** Metabolic pathway converting L-tyrosine to pC-HSL via the intermediates 4-coumaric acid and 4-coumaroyl-CoA. **C)** Schematic of the bacterial reporter circuit used to determine pC-HSL concentrations in engineered bacteria supernatant. The RpaR transcription factor activates GFP expression from an RpaO promoter in response to intracellular pC-HSL. By applying solutions containing pC-HSL and measuring GFP fluorescence, the circuit can be used to back-calculate pC-HSL levels. **D)** Quantification of pC-HSL accumulation in culture supernatants of engineered *E. coli* and *P. putida* grown with (+) or without (–) p-coumarate precursor over 48 h. Points represent the mean of n = 3 biological replicates, and the shaded regions represent the standard deviation.

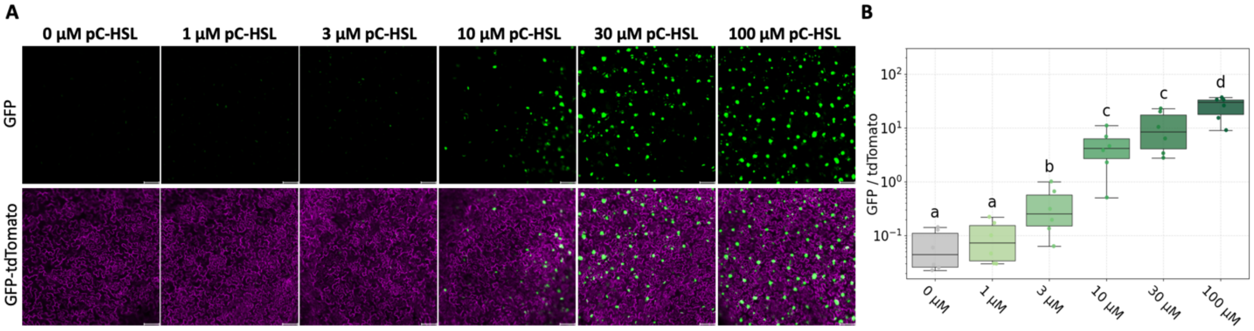


**Supplementary Figure 12. Characterization of pC-HSL-inducible circuit in sentinel plant leaves grown in soil. A)** Representative confocal images of GFP (green) and tdTomato (magenta) signals in *A. thaliana* leaves grown in soil with root-treated pC-HSL at concentrations ranging from 0 to 100 µM. Scale bars, 100 µm. **B)** Quantification of circuit activation in leaves of stably transformed *A. thaliana* seedlings. Seedlings were grown on split plates filled with soil, in which pC-HSL was applied only to the lower region. Data were shown as boxplots (median, Interquartile Range IQR, and 1.5×IQR whiskers) with individual data points overlaid (n = 6 biological replicates). Log-transformed data were analyzed by one-way ANOVA with Tukey's HSD post hoc test. Different letters indicate statistically significant differences ($p < 0.05$).
